# Rescue of K_ATP_ channel nonsense mutations using anticodon-edited transfer RNAs

**DOI:** 10.64898/2026.11.20.768202

**Authors:** Yi-Ying Kuo, Zhongying Yang, Joseph J. Porter, John D. Lueck, Show-Ling Shyng

## Abstract

K_ATP_ channels composed of sulfonylurea receptor 1 (SUR1) and Kir6.2 encoded by *ABCC8* and *KCNJ11*, respectively, play a crucial role in regulating insulin secretion. Nonsense mutations that introduce premature termination codons (PTCs) in *ABCC8* and *KCNJ11* disrupt K_ATP_ channel expression, resulting in congenital hyperinsulinism (HI). Currently, there is no effective non-invasive treatment for HI caused by K_ATP_-PTCs. Anticodon-edited transfer RNAs (ACE-tRNAs), which suppress PTCs and restore full-length protein expression, have emerged as potential therapies for diseases caused by nonsense mutations. Here, we evaluate the efficacy of ACE-tRNAs to rescue the expression and function of K_ATP_-PTCs in a heterologous COS cell expression system. In five selected SUR1-PTCs, ACE-tRNAs incorporating native amino acids at the PTC sites recovered K_ATP_ channel expression and activity. ACE-tRNA^Leu^_UGA_, a highly efficient suppressor incorporating leucine at PTCs, also rescued protein expression of all five variants; however, two of the five rescued PTC variants showed impaired gating, indicating functional intolerance to Leu substitution at these positions. An exhaustive screen of the rescue effects of ACE-tRNA^Leu^ on 100 HI-associated K_ATP_-PTCs reported to date demonstrated therapeutically meaningful functional rescue of 60 out of 92 SUR1-PTC variants and 6 out of 8 Kir6.2-PTC variants. In addition, optimized ACE-tRNA^Ser^ rescues several PTCs that were unresponsive to ACE-tRNA^Leu^, further expanding the applicability of ACE-tRNA for rescuing PTC variants. Our results demonstrate that ACE-tRNAs are effective in rescuing the expression and function of a majority of K_ATP_ PTC variants and suggest ACE-tRNAs as a promising molecular therapy for HI caused by nonsense mutations.

**Significance statement:** Nonsense mutations in K_ATP_ channel genes cause congenital hyperinsulinism, a life-threatening disorder characterized by hypoglycemia resulting from dysregulated insulin secretion. These mutations introduce premature termination codons that produce truncated proteins and impair K_ATP_ channel function, contributing to disease with limited treatment options. Here, we demonstrate that anticodon-edited transfer RNAs (ACE-tRNAs) can effectively suppress these premature stop codons to restore functional K_ATP_ channels. By broadly screening 100 identified K_ATP_ channel nonsense mutations, we show that ACE-tRNAs can rescue the expression and physiological function of the majority of variants. Our findings provide a promising therapeutic strategy for congenital hyperinsulism and establish a proof-of-concept for applying ACE-tRNA technology to a broad range of genetic diseases caused by nonsense mutations.

**Graphic abstract:** Pancreatic K_ATP_ channels, composed of SUR1 and Kir6.2 subunits (encoded by *ABCC8* and *KCNJ11*, respectively), couple intracellular ATP/ADP ratios to the β-cell membrane potential to regulate insulin secretion. When glucose levels rise, an increased ATP/ADP ratio closes K_ATP_ channels, triggering β-cell depolarization, Ca²^+^ influx through voltage-gated calcium channels, and insulin release. Premature termination codon (PTC) variants in *ABCC8* or *KCNJ11* result in truncated, nonfunctional subunits and reduced surface channel expression, causing severe congenital hyperinsulinism. Anticodon-engineered tRNAs (ACE-tRNAs) suppress premature termination by recognizing stop codons and incorporating amino acids to restore full-length SUR1 and Kir6.2 protein synthesis. Our results demonstrate that ACE-tRNAs can rescue the expression and function of the majority of SUR1 and Kir6.2 PTC variants.

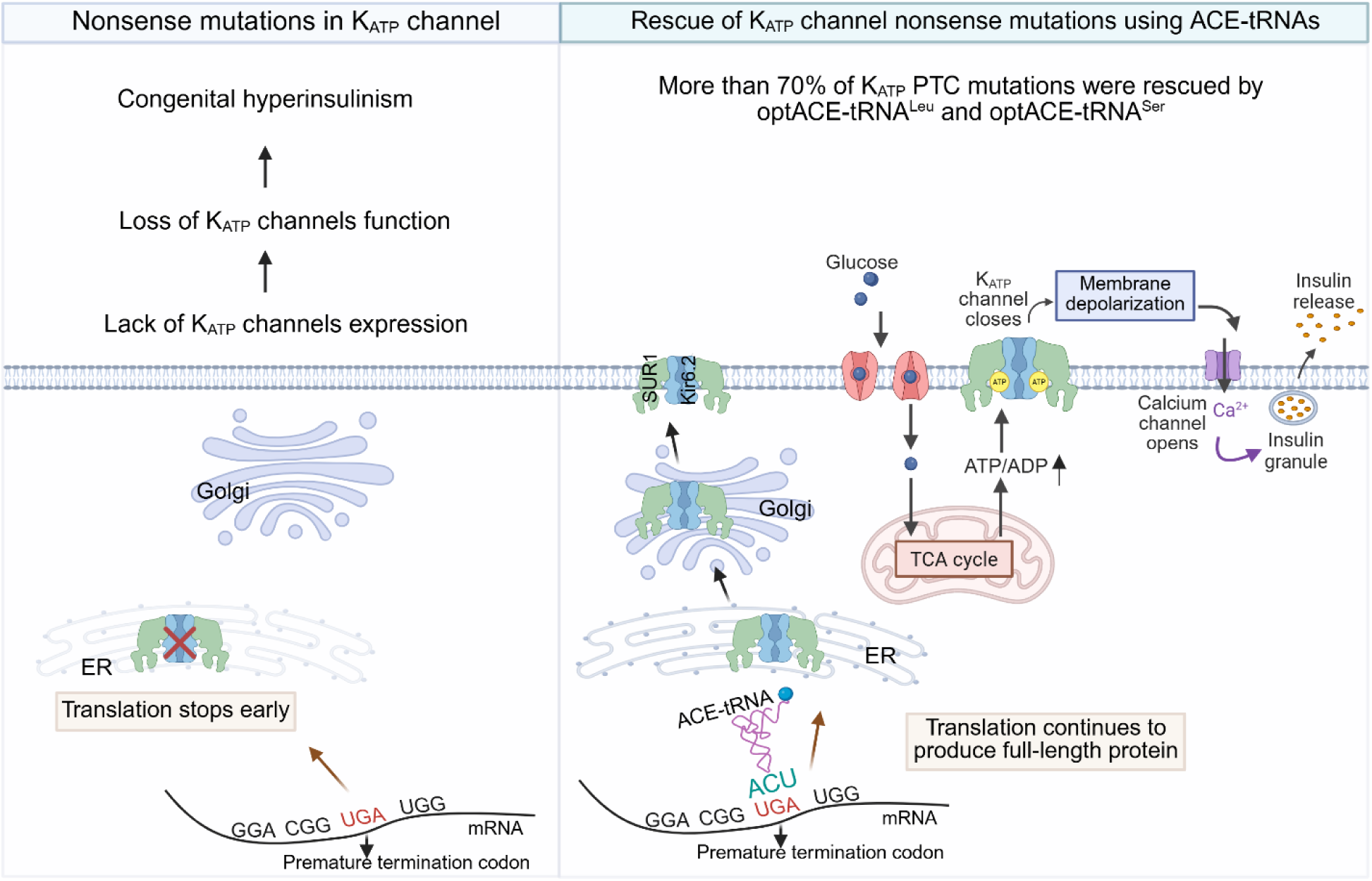

## Introduction

ATP-sensitive potassium (K_ATP_) channels in pancreatic endocrine β-cells regulate insulin secretion and are vital for glucose homeostasis (1–3). Composed of four pore-forming Kir6.2 subunits encoded by *KCNJ11*, and four regulatory sulfonylurea receptor 1 (SUR1) subunits encoded by *ABCC8*, K_ATP_ channels are gated by intracellular ATP and ADP via allosteric binding sites in the Kir6.2 and SUR1 subunits. This enables them to couple changes in the intracellular ATP-to-ADP ratio to the β-cell membrane potential. When blood glucose levels rise, an increase in the intracellular ATP-to-ADP ratio leads to K_ATP_ channel closure and membrane depolarization. This opens voltage-gated calcium channels, allowing Ca^2+^ influx to trigger insulin release (4, 5). Conversely, when blood glucose levels are low, a decrease in the ATP-to-ADP ratio opens K_ATP_ channels, maintaining β-cell membrane in a hyperpolarized resting state and preventing insulin secretion.

Genetic variants in *KCNJ11 and ABCC8* that cause complete or partial loss of K_ATP_ channel expression or function can lead to congenital hyperinsulinism (HI) (6–8), a rare disease characterized by persistent insulin secretion despite low blood glucose levels, resulting in potentially life-threatening hypoglycemia (7, 9, 10). The disease typically manifests during the neonatal period, and timely treatment is critical to prevent hypoglycemia-induced brain damage or death. However, current treatment options remain limited. Diazoxide, a K_ATP_ channel activator and the only FDA-approved oral therapy for HI (7, 11), is effective only in patients with sufficient functional K_ATP_ channels in the plasma membrane. For patients who do not respond to diazoxide, pancreatectomy is often required to manage severe hypoglycemia, which can lead to pancreatic enzyme deficiency and the development of insulin-dependent diabetes (12).

Nonsense mutations are single-nucleotide alterations in coding regions that convert a canonical amino acid codon to one of the three stop codons: TAG, TGA, or TAA (UAG, UGA and UAA respectively, in RNA) (13). The premature termination codons (PTCs) thus introduced generate truncated nonfunctional or dysfunctional proteins by terminating translation prematurely. Furthermore, nonsense mutations promote degradation of mRNAs through nonsense-mediated mRNA decay (14, 15). These mechanisms contribute to loss of expression and function of the affected protein. Several therapeutic approaches have been investigated for diseases caused by PTCs. For example, small molecules such as aminoglycosides or oxadiazoles promote PTC readthrough; however, they often suffer from low efficacy and nephrotoxicity and ototoxicity (16). Gene replacement and CRISPR-based gene editing therapies can directly correct nonsense mutations, but their broader applications face significant challenges. Gene replacement, or complementation, is constrained by vector capacity and difficult to implement for large genes such as *ABCC8*. CRISPR-based editing has potential off-target effects and requires custom design for each mutation, limiting its scalability across diverse PTC variants.

Anticodon-edited transfer RNAs (ACE-tRNAs or suppressor-tRNAs) are edited tRNAs whose anticodons are modified to base-pair with one of the three stop codons (17). ACE-tRNAs are charged with their cognate amino acids and are recognized by the endogenous translation machinery to incorporate a charged amino acid at the PTC site, allowing the translation of a full-length protein from a PTC-containing mRNA to restore protein expression and function (17, 18). There is a growing number of reports on the efficacy of ACE-tRNAs in rescuing the expression and function of PTC variants associated with disease. These include CFTR (Cystic Fibrosis Transmembrane Conductance Regulator)-PTCs associated with cystic fibrosis, CDKL5-PTCs associated with CDKL5 deficiency disorders, TCTP-PTCs associated with Batten disease, β-hexosaminidase A-PTCs associated with Tay-Sachs disease, and NPC intracellular cholesterol transporter-PTCs associated with Niemann-Pick disease type C1 (14, 19–21), suggesting the broad therapeutic applicability of ACE-tRNAs.

In K_ATP_-HI, approximately 100 nonsense mutations have been identified in *ABCC8* and *KCNJ11* (22–26) (Table S1, S2; Figure S1). These mutations, mostly in *ABCC8* coding for the large regulatory SUR1 protein, account for ∼12% of all genetic variants identified in K_ATP_-HI and are associated with diazoxide-unresponsive disease due to the lack of channel expression. The potential of ACE-tRNAs to restore the expression and function of HI-causing K_ATP_ channel PTC variants has not been determined. Here, we evaluated the applicability of ACE-tRNAs in rescuing the expression and function of K_ATP_ channels of all K_ATP_-PTC reported to date in transiently transfected COSm6 cells (experimental schematics shown in Figure S2). Our findings demonstrate that ACE-tRNAs can restore expression and function of K_ATP_ channels, supporting ACE-tRNAs as a promising therapeutic approach for HI-causing K_ATP_ channel nonsense mutations.

## Results

### Rescue of SUR1-PTC variants using ACE-tRNAs that incorporate native amino acids

To test the feasibility of K_ATP_-PTC rescue by ACE-tRNAs, we began with five *ABCC8* nonsense mutations that result in PTCs across different regions of the SUR1 protein, including R248X, R705X, R934X, C1150X, and R1539X (Figure S1; Table S1). These PTCs all have the opal stop codon TGA. Therefore, ACE-tRNAs that recognize the UGA stop codon in mRNAs and are charged with Arginine or Cysteine (denoted 4XACE-tRNA^Arg^_UGA_ or 4XACE-tRNA^Cys^_UGA_, where the superscript indicates the amino acid charged on the tRNA and the subscript indicates the stop codon recognized by the tRNA), which incorporate the endogenous amino acid (Arg or Cys) at the PTC sites, were chosen for our initial experiments.

#### Characterization of SUR1-PTCs

We first confirmed that the various SUR1-PTCs result in truncated SUR1 products. For this, N-terminally FLAG-epitope tagged SUR1 (FLAG-SUR1)-PTC constructs were transfected into COSm6 cells and protein products probed using an anti-FLAG antibody. WT FLAG-SUR1 exhibited a strong signal at ∼150 kD, predicted for full-length FLAG-SUR1. By contrast, no full-length FLAG-SUR1 signal was observed for any of the five SUR1-PTCs. Instead, truncated FLAG-SUR1 proteins at molecular masses predicted based on the positions of the PTCs were observed for R934X, C1150X, and R1539X (Figure S3). Note, no FLAG signal was detected for R248X or R705X (Figure S3), which could be due to nonsense-mediated decay of the mRNAs, resulting in low abundance of the truncated protein products. These results confirm that the introduced PTCs effectively abort protein translation at the PTCs.

Truncated SUR1 or Kir6.2 proteins resulting from K_ATP_-PTCs that are unable to assemble into channel complexes are presumably targeted for degradation (27, 28). However, whether PTCs may generate truncated proteins that retain the ability to assemble into octameric 4SUR1:4Kir6.2 K_ATP_ complexes and traffic to the plasma membrane is unknown. We tested this possibility by examining SUR1-R1539X as it results in the smallest truncation of 44 amino acids. Electrophysiology recordings of cells transfected with SUR1-R1539X plasmid and WT Kir6.2 detected very small currents that were ATP-sensitive, indicating that R1539X-SUR1 can assemble with Kir6.2 to form K_ATP_ channels. However, these truncated channels did not respond to the physiological activator MgADP nor the pharmacological activator diazoxide (Figure S4), indicating these channels are not functional under physiological conditions. We next asked whether R1539X under heterozygous conditions could co-assemble with WT SUR1 and exert a dominant-negative effect on channel function. To test this, we co-expressed Kir6.2 with a 1:1 mixture of WT SUR1 and the R1539X variant and performed electrophysiology on channels expressed in the plasma membrane. We found that under this simulated heterozygous condition, the channels showed WT-like MgADP and diazoxide responses (Figure S4), indicating that the R1539X-SUR1 does not impose a dominant-negative effect. These results led us to conclude that although SUR1-PTC products, at least R1539X, can form channels, they have negligible effects on channel expression or function. Thus, any residual K_ATP_-PTC products are unlikely to confound data interpretations in the ACE-tRNA rescue experiments.

#### Full-length SUR1 protein expression rescue by ACE-tRNAs

To determine whether ACE-tRNAs suppress PTCs to yield full-length SUR1 protein, COSm6 cells were transiently transfected with each of the five SUR1-PTC constructs and wild-type (WT) Kir6.2, along with ACE-tRNA^Sc^ (scrambled control), ACE-tRNA^Arg^_UGA_ or ACE-tRNA^Cys^_UGA,_ and tested for channel protein expression and function. Cells were harvested and subjected to immunoblotting using an antibody recognizing the C-terminal 13 amino acids of SUR1 such that only full-length SUR1 is detected (see Materials and Methods). In cells co-transfected with WT SUR1, WT Kir6.2, and ACE-tRNA^Sc^, a strong lower band and a weaker upper band corresponding to the immature core-glycosylated and mature complex-glycosylated full-length SUR1 were detected as expected. The latter represents SUR1 that has successfully co-assembled with Kir6.2 and passed the ER quality control to traffic to medial Golgi where the two N-linked glycosylation sites are modified to acquire complex-glycosylation. In cells co-transfected with SUR1-PTCs plus Kir6.2 and ACE-tRNA^Sc^, no SUR1 protein signal was detected, consistent with the SUR1-PTCs generating truncated SUR1 products that do not contain the SUR1 C-terminal epitope. In contrast, in cells co-transfected with R248X, R705X, R934X, or R1539X and ACE-tRNA^Arg^_UGA_, and C1150X and ACE-tRNA^Cys^_UGA_, full-length core-glycosylated SUR1 was clearly observed, with complex-glycosylated SUR1 also visible (Figure 1*A*).

**Figure 1.**
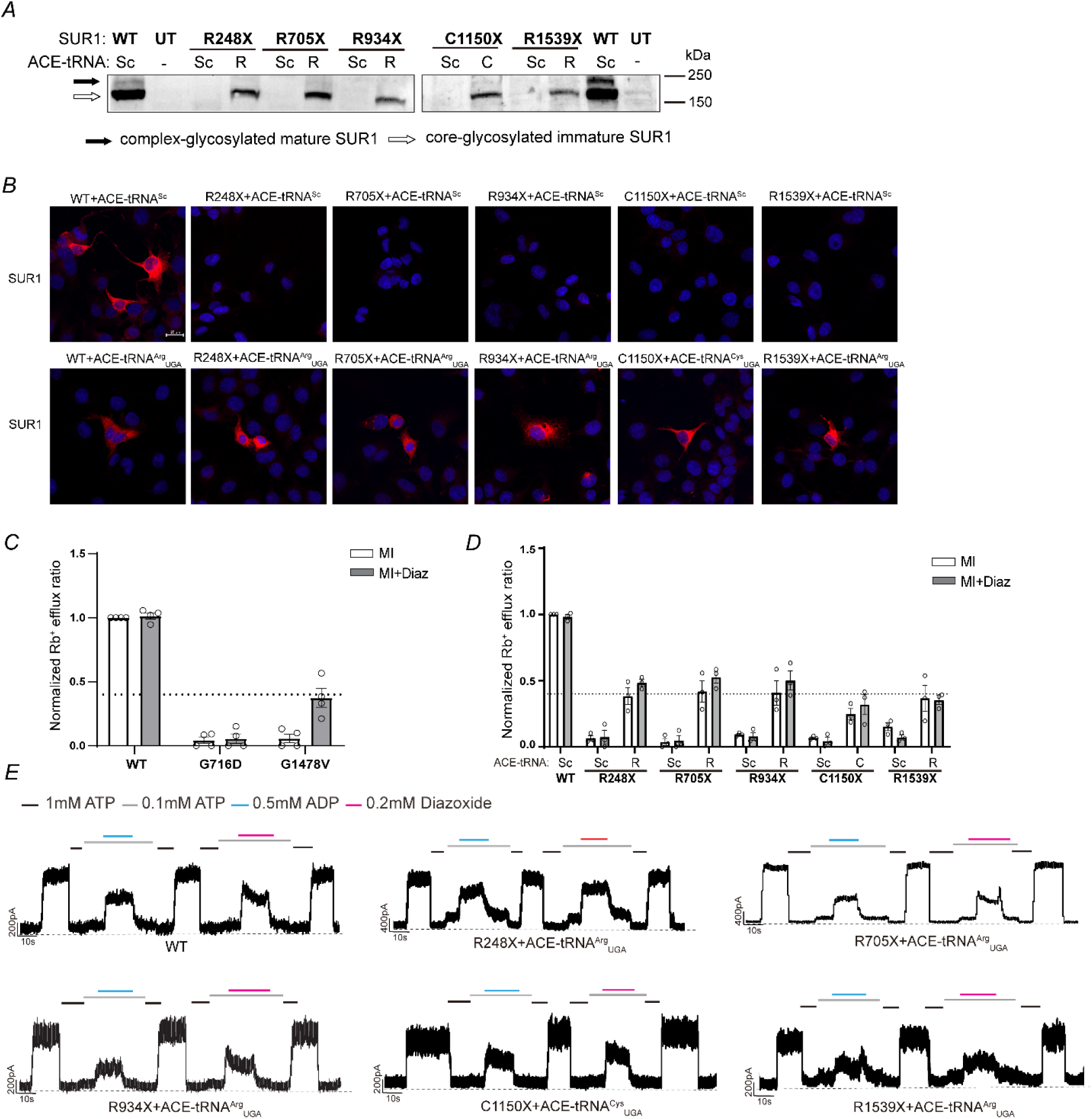
ACE-tRNA^Arg^_UGA_ and ACE-tRNA^Cys^_UGA_ rescue Arginine and Cysteine SUR1-PTC variants by incorporating the native amino acid. (*A)* Immunoblots show addition of ACE-tRNAs, which incorporate native amino acids at the PTC sites, results in the expression of full-length SUR1 protein. (*B)* Representative immunofluorescence images show SUR1 expression (in red) in COSm6 cells transfected with R248X, R705X, R934X, C1150X and R1539X with ACE-tRNA^Sc^, ACE-tRNA^Arg^_UGA_ or ACE-tRNA^Cys^_UGA_ (scale bar is 20 μm). (*C)* A diazoxide-unresponsive mutation (G716D) and a diazoxide-responsive mutation (G1478V) were evaluated under MI and MI+Diaz conditions. Based on G1478V results, an Rb^+^ efflux ratio of 40% relative to WT was set as the therapeutic threshold required for effective channel rescue. (*D)* Rb^+^ efflux assay results show the rescue effects of ACE-tRNA^Arg^_UGA_ and ACE-tRNA^Cys^_UGA_ on SUR1-PTC variants under metabolic inhibition (MI) and metabolic inhibition plus diazoxide (MI+Diaz). The dashed line marks 40% of WT therapeutic threshold. (Data are presented as means ± SEM) (*E)*. Patch-clamp recordings demonstrate that ACE-tRNA-rescued SUR1-PTC channels respond to MgADP and diazoxide like WT channels (holding potential: −50 mV; upward deflections indicate inward currents; black dashed lines indicate the 0 current baseline in 1 mM MgATP).

Expression of full-length SUR1 was further corroborated by immunofluorescence staining experiments using an anti-SUR1 C-terminal peptide antibody. Fluorescence signals for SUR1 were observed in WT-SUR1 expressing cells and cells co-transfected with SUR1-PTCs and their cognate ACE-tRNAs. In contrast, no signals were detected in control cells transfected with SUR1-PTCs and ACE-tRNA^Sc^ (Figure 1*B*). Together, these results demonstrate that ACE-tRNA-dependent incorporation of native amino acids at SUR1-PTCs efficiently restore full-length SUR1 expression.

#### Functional rescue of K_ATP_ channels containing SUR1-PTCs

To evaluate the function of K_ATP_ channels encoded from transcripts containing PTCs that are suppressed by ACE-tRNAs, we employed Rb^+^ efflux assays and electrophysiology (Figure *1D, E*). The Rb^+^ efflux assay uses Rb^+^ as a surrogate of K^+^ to monitor K_ATP_ channel activity in intact cells (29) (see Materials and Methods and Figure S2*B*). Cells transfected with PTC variants or WT channels were treated with metabolic inhibitors (MI; 1 mM deoxyglucose and 2.5 µg/ml oligomycin) immediately prior to the assay, which lowered the intracellular ATP to ADP ratio to open K_ATP_ channels. In a second series of experiments, cells were treated with MI along with diazoxide at 200 µM (MI+Diaz). Diazoxide is a K_ATP_ channel opener clinically used to treat HI and may further potentiate K_ATP_ channel activity. The Rb^+^ efflux ratio (efflux during a 30 min period relative to total Rb^+^ at time 0; see Materials and Methods) observed in cells transfected with the PTC variants was normalized to that in cells transfected with WT channel control to assess K_ATP_ channel activity.

In cells co-transfected with various SUR1-PTC variant channels and the ACE-tRNA^Sc^ control, there was little Rb^+^ efflux under both MI and MI+Diaz conditions consistent with lack of channel expression. In contrast, cells co-transfected with the SUR1-PTC variants and their cognate ACE-tRNAs all showed significant increase in Rb^+^ efflux under both MI and MI+Diaz conditions, with diazoxide showing small potentiation effects for most mutations (Figure 1*D*). Note for WT channels, MI already resulted in maximum Rb^+^ efflux under our experimental condition such that addition of diazoxide had little potentiation effects. To establish the therapeutic benchmark of ACE-tRNAs rescue in the Rb^+^ efflux assay, we also examined two non-PTC SUR1 mutations, G716D and G1478V, which have been associated with diazoxide unresponsive and responsive HI, respectively (30). The diazoxide-unresponsive mutation G716D showed little Rb^+^ efflux under both MI and MI+Diaz conditions, similarly to PTC mutations in the absence of ACE-tRNA rescue. Conversely, the diazoxide-responsive mutation G1478V, while showing very low efflux in MI, exhibited an Rb^+^ efflux ratio of approximately 40% (0.37±0.07) that of WT in MI+Diaz (Figure 1*C*). The G1478V mutation is one of the HI mutations previously shown to respond poorly to decreased ATP/ADP ratios but still retain sufficient activity to be clinically responsive to diazoxide treatment (30). Therefore, the Rb^+^ efflux ratio of G1478V of ∼ 40% of WT observed in MI+Diaz was taken as the therapeutic threshold for ACE-tRNAs rescue (indicated by a dashed line shown in all Rb efflux figures). Based on this criterion, the rescue of R248X, R705X and R934X by ACE-tRNA^Arg^_UGA_ was all at or above the 40% WT activity threshold. Under MI, the corresponding Rb^+^ efflux ratios were 0.38 ± 0.06, 0.42 ± 0.08, and 0.41 ± 0.09, respectively. Under MI+Diaz condition, the corresponding ratios were 0.48 ± 0.03, 0.53 ± 0.04, and 0.50 ± 0.07. R1539X and C1150X were rescued by ACE-tRNA^Cys^_UGA_ being lower but nevertheless approaching 40% of the WT (Figure *1D*). The Rb^+^ efflux ratios for R1539X under MI and MI+Diaz conditions were 0.36 ± 0.10 and 0.35 ± 0.03, respectively, whereas those for C1150X were 0.25 ± 0.04 and 0.32 ± 0.08, respectively.

Restored PTC variant channels were further tested for their gating properties using inside-out patch-clamp electrophysiology recordings (see Materials and Methods and Figure S2*C*). Specifically, we focused on channel responses to stimulation by the physiological activator MgADP and pharmacological activator diazoxide, both frequently impaired in HI-associated K_ATP_ variants. In all cases, restoration of full-length channels by cognate ACE-tRNAs showed MgADP and diazoxide responses indistinguishable from WT channels (Figure 1*E*), reflecting the accuracy of cognate ACE-tRNAs in incorporating native amino acids at the PTC sites. Collectively, the Rb^+^ efflux and electrophysiology results demonstrate that ACE-tRNA^Arg^_UGA_ and ACE-tRNA^Cys^_UGA_ significantly increased the expression and function of K_ATP_ channels containing SUR1-R248X, R705X, R934X, C1150X, and R1539X PTCs.

### Assessing the efficacy of ACE-tRNA^Leu^_UGA_ to rescue SUR1-PTCs

Functional rescue of SUR1-PTCs by ACE-tRNAs is dependent on both the readthrough efficiency of the PTCs by an ACE-tRNA and the functional impact of the amino acid the ACE-tRNA incorporates at the PTC site. Of the ACE-tRNAs reported, ACE-tRNA^Leu^_UGA_ shows superior PTC readthrough efficiency (14). We wondered whether the above five SUR1-PTC variants could be rescued by ACE-tRNA^Leu^_UGA_, which incorporates leucine rather than the native amino acid at the PTC sites. To test this, COSm6 cells were co-transfected with the SUR1-PTC construct and WT Kir6.2 along with an optimized ACE-tRNA^Leu^_UGA_ (opt.ACE-tRNA^Leu^_UGA_), which was recently reported to have further improved PTC suppression efficiency as compared to the first-generation ACE-tRNA^Leu^_UGA_ (31). Immunoblotting results showed that opt.ACE-tRNA^Leu^_UGA_ led to the appearance of full-length SUR1 in all five PTC variants to varying degrees (Figure 2*A*). These results demonstrate the capacity of ACE-tRNA^Leu^_UGA_ to rescue full-length SUR1 expression in all five PTCs. Functional testing using Rb^+^ efflux assays showed that rescue of R248X, R705X, R934X, and C1150X by opt.ACE-tRNA^Leu^_UGA_, restored efflux to variable degrees well above 40% of WT levels in both MI and MI+Diaz conditions; in contrast, R1539X showed no or poor efflux response to MI or MI+Diaz, well below 40% of WT channel activity (Figure 2*B*).

**Figure 2.**
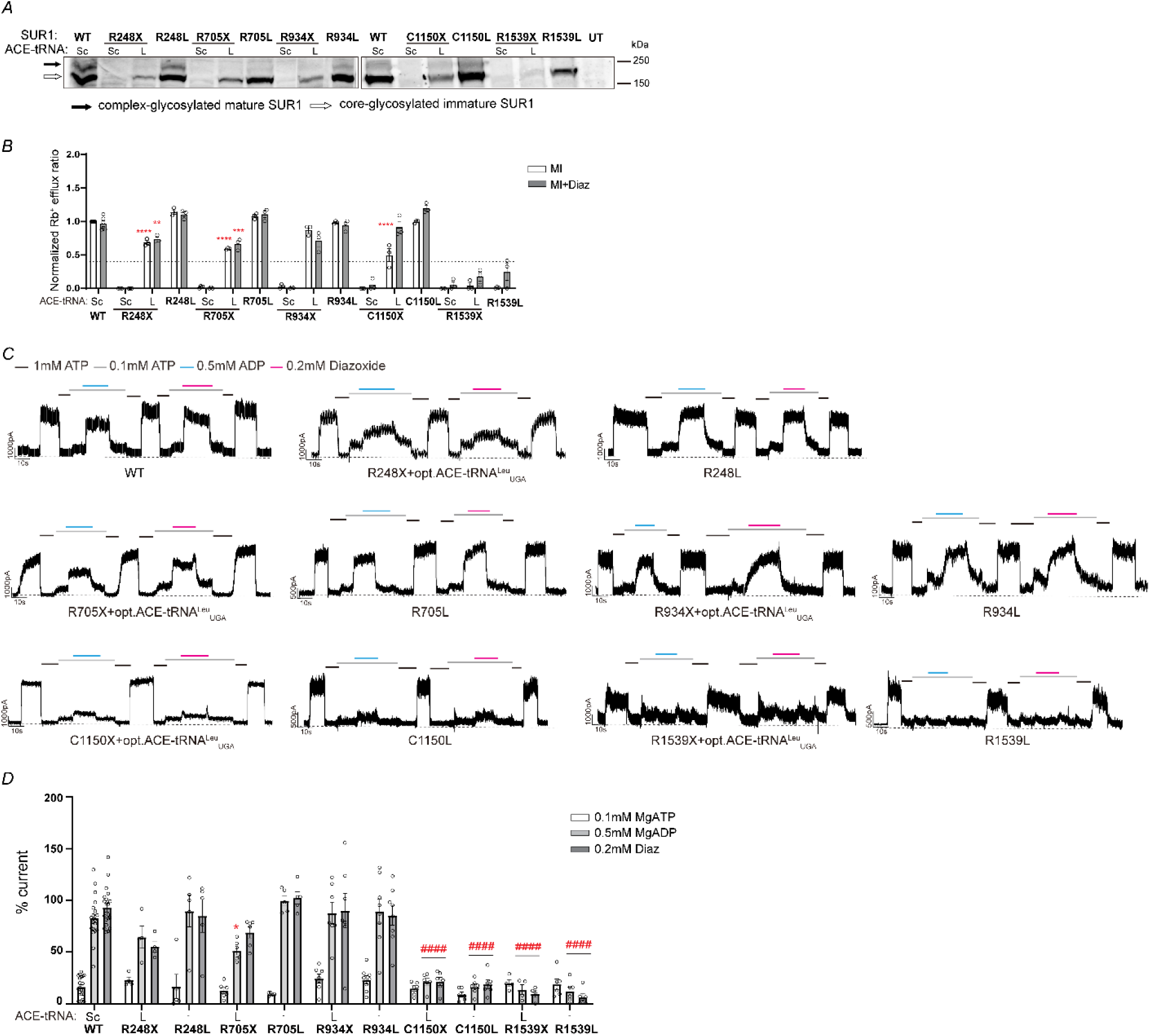
Efficacy of optimized ACE-tRNA^Leu^_UGA_ in rescuing the expression and function of SUR1 PTCs. (*A)* Immunoblots showing SUR1 full-length protein expression levels in cells co-transfected with SUR1-PTC R248X, R705X, R934X, C1150X, or R1539X and WT Kir6.2, along with scrambled control plasmid (Sc) or plasmid encoding the optimized ACE-tRNA^Leu^_UGA_ (L). Lysates from cells co-transfected with WT Kir6.2 and SUR1 with direct leucine substitution R248L, R705L, R934L, C1150L, or R1539L were run side-by-side as controls. Core-glycosylated SUR1 is indicated by the empty arrow and the complex-glycosylated SUR1 by the solid arrow. (*B)* Rb^+^ efflux assay results of opt.ACE-tRNA^Leu^_UGA_-rescued PTCs and leucine substitution controls. Data are presented as means ± SEM. Statistical significance was assessed at α = 0.05 using one-way ANOVA followed by Tukey’s multiple comparisons test, with MI and MI+Diaz responses analyzed separately. Statistical significance comparing opt.ACE-tRNA^Leu^_UGA_-rescued PTCs and direct leucine substitutions under MI or MI+Diaz is indicated as follows: \*\*\*\**p* < 0.0001, \*\**p* < 0.01. (*C)* Representative electrophysiology traces of the opt.ACE-tRNA^Leu^_UGA_-treated and leucine substitution controls (holding potential: −50 mV; upward deflections indicate inward currents; black dashed lines indicate the 0-current baseline in 1 mM MgATP). (*D)* Quantification of MgADP and diazoxide responses in (*C)*. Data are presented as means ± SEM. Statistical significance was assessed at α = 0.05 using one-way ANOVA followed by Tukey’s multiple comparisons test, with MgADP and diazoxide responses analyzed separately. Statistical significance between optACE-tRNA^Leu^_UGA_-rescued channels and leucine substitution control: \**p* < 0.05, Statistical significance compared to WT is indicated as follows: ^####^*p* < 0.0001, ^#^*p* < 0.05.

As a control, we evaluated the impact of direct Leu substitution at the five SUR1-PTC sites via site-directed mutagenesis on channel expression and function. Western blots showed that all five leucine substitution mutations: R248L, R705L, R934L, C1150L, and R1539L, were tolerated by SUR1, although R1539L was expressed to a lower extent than the others (Figure 2*A*). In Rb^+^ efflux assays, the R248L, R705L, R934L, and C1150L variants all displayed WT-channel like efflux in both MI and MI+Diaz conditions. In contrast, the R1539L variant showed no efflux in MI and very low activity in MI+Diaz, similar to that observed for opt.ACE-tRNA^Leu^_UGA_-rescued R1539X (Figure 2*B*).

For R248X, R705X, R934X, and C1150X, the lower SUR1 protein levels seen in western blots and lower channel activity in Rb^+^ efflux assays upon rescue by opt.ACE-tRNA^Leu^_UGA_-compared to their corresponding direct leucine substitution mutants suggest that while PTC suppression by opt.ACE-tRNA^Leu^_UGA_ is generally efficient, it remains incomplete. For R1539X, the lack of efflux activity observed upon opt.ACE-tRNA^Leu^_UGA_ rescue as well as in the R1539L suggests leucine substitution at this site has adverse effects on the channel function.

To investigate whether the varied functional rescue observed in Rb^+^ efflux assays arose from altered channel gating properties, we first performed inside-out patch-clamp recording experiments to test nucleotide and diazoxide sensitivity of SUR1-PTC channels in cells co-expressing opt.ACE-tRNA^Leu^_UGA_. All five variants showed pronounced currents in K-INT solution, indicating the presence of K_ATP_ channels in the plasma membrane. However, while the opt.ACE-tRNA^Leu^_UGA_-rescued R934X channels had MgADP and diazoxide responses comparable to WT channels, rescued R248X and R705X channels showed a trend of slightly reduced MgADP and diazoxide response which did not reach statistical significance, and rescued C1150X and R1539X channels showed dramatically impaired response to both MgADP and diazoxide, with R1539X channels showing the most severe impairment (Figure 2*C*). Of note, despite the markedly reduced MgADP and diazoxide response in electrophysiology experiments, the C1150X channels still exhibited significant activity in Rb^+^ efflux experiments. This discrepancy is likely due to the different designs of the two experiments: electrophysiology records real-time channel activity, whereas the Rb^+^ efflux assay measures the accumulated release of Rb^+^ over a 30-min period. Thus, even though the C1150X channels exhibited markedly reduced MgADP and diazoxide responses in electrophysiological recordings, the residual channel responses were sufficient to produce significant Rb^+^ efflux over time. In contrast, the opt.ACE-tRNA^Leu^_UGA_-rescued R1539X channels failed to be stimulated by MgADP or diazoxide in electrophysiology experiments. In fact, currents in 0.1 mM MgATP plus 0.5 mM MgADP or plus 0.2 mM diazoxide were even smaller than currents observed in 0.1 mM MgATP (Figure 2*D*). These observations are consistent with opt.ACE-tRNA^Leu^_UGA_-rescued R1539X channels showing background or very low Rb^+^ efflux activity, which is consistent with the channel not being opened in response to metabolic inhibition and diazoxide.

Parallel electrophysiology recordings of the corresponding five Leu substitution mutants R248L, R705L, R934L, C1150L, and R1539L were carried out as controls. Results from the five Leu substitution mutant generally followed the trends of those obtained from opt.ACE-tRNA^Leu^_UGA_-rescued R248X, R705X, R934X, C1150X, and R1539X, with R248L, R705L, and R934L being close to WT, while C1150L and R1539L showed greatly impaired MgADP and diazoxide responses. Thus, while leucine substitution at the R248, R705, and R934 positions is well tolerated, it is not tolerated at the C1150 and R1539 positions for preserving WT channel gating response to MgADP and diazoxide. Of note, the MgADP and diazoxide responses of opt.ACE-tRNA^Leu^_UGA_-rescued R248X and R705X appeared smaller than their R248L and R705L counterparts, with the difference in MgADP response between opt.ACE-tRNA^Leu^_UGA_-rescued R705X and the R705L reaching statistical significance. The cause of this small difference is unclear but could arise from co-assembly of residual truncated R705X SUR1 with full-length R705X SUR1 rescued by opt.ACE-tRNA^Leu^_UGA_, although this effect is expected to be negligible based on our analysis of another PTC, SUR1-R1539X (see Figure S4). Regardless, the electrophysiology results reveal the SUR1-PTC site-specific functional consequences upon ACE-tRNA^Leu^_UGA_-mediated PTC suppression and highlight the need for experimental testing to identify PTCs suitable for rescue by ACE-tRNA^Leu^ as a platform approach.

### Screening of SUR1 and Kir6.2 PTCs for optACE-tRNA^Leu^ and optACE-tRNA^Ser^ suppression

Having established the efficacy of optACE-tRNA^Leu^ to rescue the expression of the aforementioned-five SUR1-PTCs and the resulting channel function in some of them, we next tested the broader applicability of optACE-tRNA^Leu^ on all K_ATP_ channel PTCs reported to date. In total, there are 92 SUR1 PTC variants including the five described above (Figures 1, S1 and Table S1), and 8 Kir6.2 PTC variants (Figure S1 and Table S2) with TGA, TAG or TAA stop codons. Depending on the stop codon at the PTC site, opt.ACE-tRNA^Leu^_UGA_, opt.ACE-tRNA^Leu^_UAG_ or opt.ACE-tRNA^Leu^_UAA_ were used. Full-length SUR1 and Kir6.2 protein expressions were confirmed by immunoblotting for all SUR1 and Kir6.2 PTCs (Figure S5). The Rb^+^ efflux assay results showed that 60 out of 92 SUR1-PTC variants had efflux ratios higher than 40% of the WT under either MI or MI+Diaz condition (Figure 3*A*). Additionally, 6 out of 8 Kir6.2 PTC variants exhibited an efflux ratio higher than 40% of the WT (Figure 3*B*).

**Figure 3.**
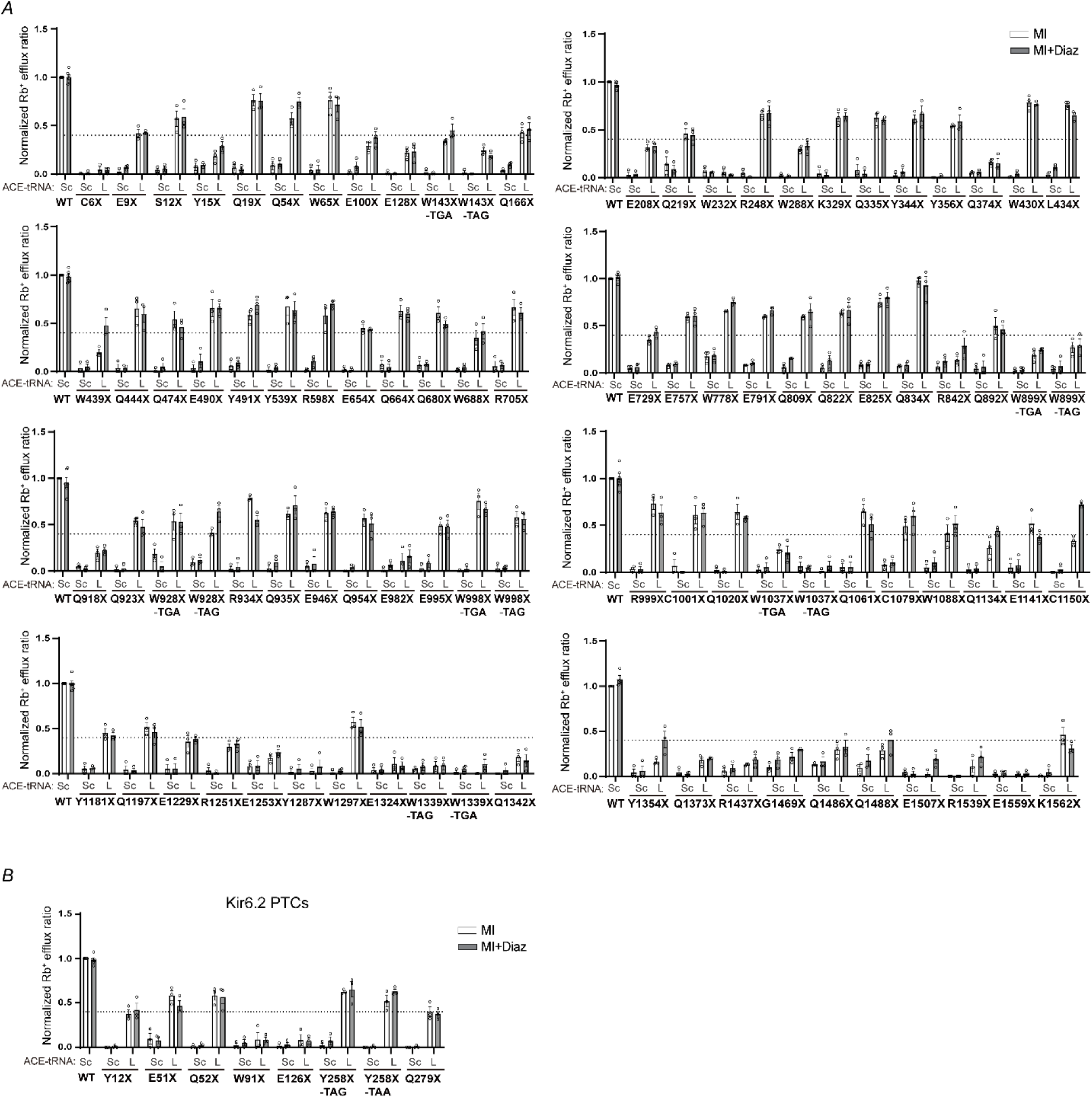
Rescue effect of opt.ACE-tRNA^Leu^ on K_ATP_-PTC variants assessed by Rb^+^ efflux assays under MI and MI+Diaz conditions. (*A)* Ninety-two SUR1 PTC mutations were tested, of which sixty showed efflux activity above or at 40% that of WT channels in the presence of opt.ACE-tRNA^Leu^. (*B)* Eight Kir6.2 PTC variants were examined, of which six showed efflux activity above or at 40% that of WT channels in the presence of opt.ACE-tRNA^Leu^.

Since leucine is a nonpolar, hydrophobic amino acid, which may not be tolerated at some sites as demonstrated for C1150X and R1539X, we also tested opt.ACE-tRNA^Ser^, which incorporates a polar, hydrophilic serine for those PTCs that failed to show efflux ratios above 40% of WT upon opt.ACE-tRNA^Leu^ rescue. Recent studies showed that opt.ACE-tRNA^Ser^ also has high PTC suppression efficiency, even surpassing opt.ACE-tRNA^Leu^ (31). We found that 6 out of 31 SUR1-PTC variants exhibited efflux ratios higher than 40% of the WT under either MI or MI+Diaz conditions (Figure 4A). However, the two Kir6.2 PTC variants did not exhibit an efflux ratio greater than 40% of the WT (Figure 4*B*). Together, our comprehensive screen identified more than 70% of K_ATP_ PTC mutations that were rescued by opt.ACE-tRNA^Leu^ and opt.ACE-tRNA^Ser^ using 40% WT channel efflux level as a therapeutic threshold, highlighting the general applicability of the opt.ACE-tRNA^Leu^ and opt.ACE-tRNA^Ser^ platforms for restoring K_ATP_ channel function by efficient suppression of nonsense mutations.

**Figure 4.**
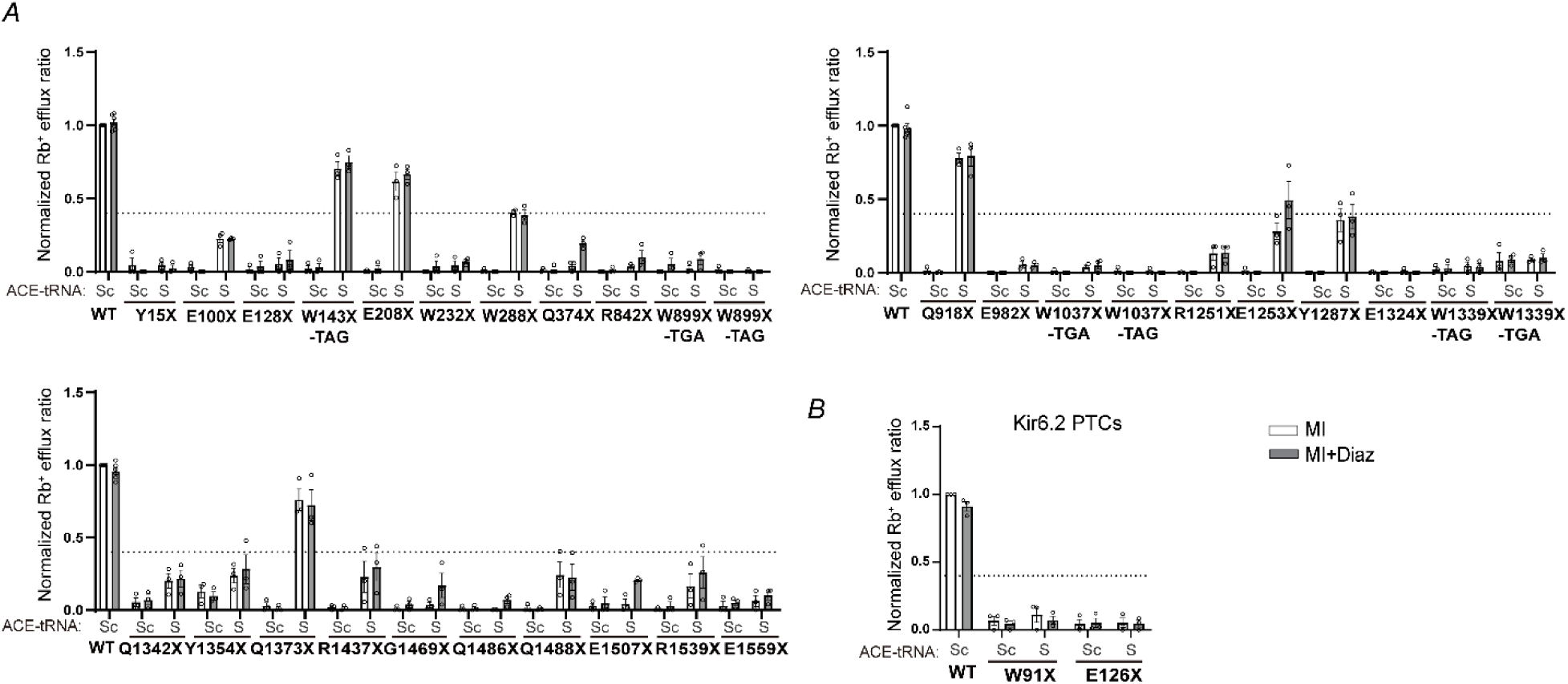
Rb^+^ efflux assays testing the rescuing effect of an optimized ACE-tRNA^Ser^ on the function of K_ATP_-PTC variants. (*A)* 31 SUR1 PTC variants were tested, of which 6 were rescued by opt.ACE-tRNA^Ser^. (*B)* 2 Kir6.2 PTC variants were examined, neither of which were rescued by opt.ACE-tRNA^Ser^.

## Discussion

Nonsense mutations are prevalent causes of human disease, accounting for 24% of pathogenic alleles (32), and remain challenging to treat. Our current study provides proof-of-concept that ACE-tRNAs suppress congenital hyperinsulinism-causing nonsense mutations in genes encoding SUR1 and Kir6.2 subunits of the K_ATP_ channel complex, thereby restoring channel protein expression and, in most cases, channel function to levels expected to be therapeutically significant, based on an Rb^+^ efflux threshold determined using a previously identified diazoxide-responsive disease mutation.

In selecting an ACE-tRNA for optimum functional restoration of K_ATP_-PTCs, two important factors should be considered as exemplified by our in-depth analysis of five selected SUR1 nonsense mutations. First, while an ACE-tRNA charged with a native amino acid at a given PTC site would preserve the function of the rescued full-length protein, in this case the channel, ACE-tRNAs charged with all amino acids are not available. Moreover, existing ACE-tRNAs charged with different amino acids have different PTC suppression efficiencies, for instance ACE-tRNA^Trp^_UGA_ displays lower efficiency than ACE-tRNA^Leu^_UGA_ (14, 17, 19, 31). Second, although ACE-tRNA^Leu^ displays more efficient nonsense suppression as compared to other ACE-tRNAs, it may not work for PTCs that do not tolerate Leu for protein folding, assembly, and/or function (33). This is well illustrated here with SUR1-C1150X and R1539X. Incorporation of Leu at these sites severely compromised the ability of the resulting K_ATP_ channels to respond to the physiological activator MgADP and the pharmacological activator diazoxide, making ACE-tRNA^Leu^ a poor choice for functional rescue of these specific K_ATP_-PTCs. Successful rescue of PTCs using ACE-tRNAs therefore depends on not only the suppression efficiency but also the channel’s tolerance of a specific amino acid at the PTC site.

It is not always possible to know *a priori* whether a given PTC site can tolerate an amino acid substitution. Indeed, comparisons between the functional results of ACE-tRNA^Leu^- or ACE-tRNA^Ser^-rescued K_ATP_ channels determined by Rb^+^ efflux assays and the predicted pathogenic effects of leucine or serine substitutions at the PTC sites using the artificial intelligence-based platform AlphaMissense (34) revealed little correlation between the experimental findings and predicted effects (Figure S6). However, incompatible amino acid substitutions can be expected in some cases. For example, C6 in SUR1 forms a disulfide bond with C26 that is critical for protein folding, trafficking and structural stability. Previous studies have demonstrated that disruption of this disulfide bond results in reduced surface expressed K_ATP_ channels (35–37). Therefore, rescuing PTC variants such as C6 would require incorporation of cysteine to maintain the function of K_ATP_ channels. Incorporation of Leu, a large hydrophobic residue, at PTC sites occupied by native amino acids that have very different physical and chemical properties, may also be a predictor of compatibility, especially in structural domains known to be important for channel expression and function. An example is seen with R1539X channels rescued by ACE-tRNA^Leu^ and confirmed by the R1539L direct Leu substitution mutation, where the resulting channels fail to respond to MgADP and diazoxide (Figure 2).

Given the potential adverse effects of non-native Leu incorporation, it is remarkable that roughly two-thirds of the K_ATP_-PTC variants tested showed therapeutically meaningful rescue by ACE-tRNA^Leu^ in Rb^+^ efflux assays (≥ 40% WT activity), with many of them having polar or even charged native amino acids including serine, cysteine, glutamine, tyrosine, glutamate, and arginine. Among the 33 K_ATP_-PTCs that did not respond well to ACE-tRNA^Leu^, six showed therapeutically meaningful Rb^+^ efflux levels in response to ACE-tRNA^Ser^, which incorporates the polar serine at the PTCs (Figure 4). Importantly our thorough screen of all HI-associated K_ATP_-PTCs reported to date using ACE-tRNA^Leu^ and ACE-tRNA^Ser^ has identified the majority of K_ATP_-PTCs as promising candidates for ACE-tRNA based therapy. As the ACE-tRNA technology continues to evolve, it is envisioned that the remaining K_ATP_-PTCs intractable to ACE-tRNA^Leu^ and ACE-tRNA^Ser^ would be rescuable when more efficient ACE-tRNAs decoding all three stop codons and charged with different amino acids become available.

Interestingly, most of the SUR1-PTCs toward the 3’ end of the gene showed poor rescue responses to ACE-tRNA^Leu^ (Figure 3). While Leu intolerance for channel function at these sites could be a cause, other factors may also contribute to the poor rescue outcomes. Previous studies have shown that the PTC nucleotide context and 3’UTR length influence ACE-tRNA suppression efficiency. Readthrough efficiency has been positively correlated with 3’UTR length, defined as the number of nucleotides from PTC to the poly-A tail, suggesting that a longer 3’UTR is associated with higher readthrough efficiency (38, 39). A PTC variant located closer to the 3’ end of a transcript with a shorter 3’UTR length is thus expected to have a reduced readthrough efficiency. This effect is likely linked to poly-A-binding protein (PABP), which plays a role in translation termination. When the 3’UTR is long and further away from PABP, the translation termination process is harder to initiate, allowing ACE-tRNA to work effectively. In addition to 3’UTR length, the stop codon context affects readthrough efficiency. The efficiency is highest for UGA, followed by UAG and UAA (19, 38). Furthermore, the first letter after the stop codon (the +4 position, where the first nucleotide of the stop codon is marked as +1) also affects readthrough efficiency. For a given stop codon, the hierarchy of readthrough efficiency has been reported as C > U > A/G (38, 40, 41). For instance, although K1562X is located near the end of the *ABCC8* gene with the shortest 3’UTR of all the SUR1-PTCs and is mutated into a UAG stop codon, its nucleotide at the +4 position is a C. This specific sequence context promotes translational readthrough, resulting in the modest rescue effect observed in our data. In contrast, E1559X showed no rescue by ACE-tRNA^Leu^; in this case, the +4 position is a U after a UAG stop codon, which presents a lower readthrough efficiency context (42). Thus, nucleotide sequence contexts also need to be considered in the readthrough efficiency near the 3’ end of the transcript.

In conclusion, our study provides the first evidence that ACE-tRNAs can overcome nonsense mutations in the pancreatic K_ATP_ channel genes to restore channel expression and function. The therapeutic applicability of ACE-tRNAs has been demonstrated in several disease models. For example, AAV-mediated delivery of ACE-tRNA has been shown to rescue a mouse model of human mucopolysaccharidosis type I (MPS-I), a lysosomal storage disease caused by nonsense mutations (43). A similar study has demonstrated that the blindness in a mouse model of Leber Congenital Amaurosis type 16, caused by the *Kcnj13* nonsense mutation, was rescued by adenovirus-mediated delivery of ACE-tRNA (44). Combined with previously reported safety data and our findings, these results support ACE-tRNAs as a highly applicable and potent therapeutic strategy for a wide spectrum of diseases caused by nonsense mutations, including congenital hyperinsulinism.

## Materials and methods

### Molecular biology

In all experiments, human FLAG epitope-tagged SUR1 (referred to as f-SUR1) cDNA was cloned into the pCMV6b (generously provided by Dr. Joseph Bryan), and human Kir6.2 in pcDNA3.1 was used. The FLAG epitope (DYKDDDDK) was inserted at the N terminus of human SUR1 cDNA, and the FLAG epitope does not affect channel function; details can be found in previous publications(23, 45). PTC or point mutations were introduced into human SUR1 using the QuikChange site-directed mutagenesis kit (Stratagene). All mutant constructs were sequence-verified, and clones from two independent PCR reactions were analyzed in each experiment to minimize the risk of artifacts from undesired PCR-introduced mutations. ACE-tRNA plasmids expressing four copies (4X) of arginine (ACE-tRNA^Arg^_UGA_), cysteine (ACE-tRNA^cys^_UGA_), leucine (ACE-tRNA^leu^_UGA_), optimized ACE-tRNA constructs expressing leucine for all three stop codons (opt.ACE-tRNA^leu^_UGA_, opt.ACE-tRNA^leu^_UAA_, opt.ACE-tRNA^leu^_UAG_), and empty vector control (ACE-tRNA^Sc)^ are as described previously (17, 31).

### Transfection of COSm6 cells with recombinant DNA

COSm6 cells were plated in a 12-well tissue culture plate and transiently transfected at approximately 80% confluency. A total amount of 0.5μg of SUR1 or Kir6.2 cDNAs containing PTC variants was co-transfected with 0.5μg of wild-type Kir6.2 or SUR1 cDNAs, respectively. Additionally, either 4X ACE-tRNAs (0.5μg) or opt.ACE-tRNAs (0.1μg) were co-transfected with the mutant K_ATP_ channels using FuGENE6^®^ (Cat#: E2691; Promega) following the manufacturer’s protocol. Plasmids and FuGENE^®^ (3μl) were first pre-mixed in Opti-MEM in a separate tube, then mixed and incubated for at least 15 minutes to ensure efficient uptake of multiple plasmids required for proper K_ATP_ channel expression.

### Immunoblotting

Forty-eight hours after transfection, Cosm6 cells were harvested and lysed on ice for 30 minutes in a buffer containing 50 mM Tris·HCl (pH 7.0), 150 mM NaCl, and 1% TritonX-100, supplemented with cOmplete^TM^ protease inhibitors (Cat#: 11836170001; Roche Applied Science), followed centrifugation at 16000 ×g for 15 min at 4 ^◦^C. The supernatants were collected, and proteins were separated by 7.5 % or 12 % SDS-PAGE, then transferred onto nitrocellulose membranes. The membranes were incubated with a primary anti-SUR1 antibody, which was raised against a C-terminal peptide of SUR1 (KDSVFASFVRADK). Then, horseradish peroxidase-conjugated secondary antibodies were applied. Signals were detected using an enhanced chemiluminescence detection kit (Cat#: 32106; Thermo Fisher). α-Tubulin was used as a loading control and was also probed on the blots.

### Immunofluorescence staining

Immunofluorescence staining of SUR1 was performed 48 hours after transfection on COSm6 cells transfected with various SUR1-PTC and ACE-tRNA as shown plus Kir6.2 constructs. Cells were washed with ice-cold phosphate-buffered saline (PBS, 137 mM NaCl, 2.7 mM KCl, 10 mM Na₂HPO₄, and 1.8 mM KH₂PO₄ at pH 7.4). After washing with cold PBS, cells were fixed with 4% paraformaldehyde for 15 minutes on ice and permeabilized with 0.75% Tween-20 for 20 minutes. The fixed cells were then blocked with 2% bovine serum albumin (BSA) in PBS for 30 minutes. Cells were incubated with primary SUR1 antibody (diluted in Opti-MEM with 0.01% BSA) for 1 hour at room temperature. After washing, cells were incubated with an Alexa Fluor 546 conjugated goat anti-rabbit secondary antibody (Invitrogen; diluted 1:200 in Opti-MEM with 0.01% BSA) for 1 hour at room temperature. The nuclei were counterstained with DAPI, and coverslips were mounted onto microscope slides using Fluoromount aqueous mounting medium (Cat#: F4680, Sigma). Images were captured using a Zeiss confocal microscope (LSM 900).

### Rb^+^ efflux assays

Cosm6 cells were transfected as described above, and untransfected cells were used as background controls. The detailed experimental procedures were described by ElSheikh et al. (29). Cells were split and incubated overnight with 5.4 mM RbCl in DMEM medium (DMEM medium, 10% fetal bovine serum, 1% penicillin-streptomycin). The following day, cells were rapidly washed twice with Ringer’s solution (5.4 mM KCl, 150 mM NaCl, 1mM MgCl₂, 0.8 mM NaH₂PO₄, 2 mM CaCl₂, 25 mM HEPES, pH 7.2). Cells were then incubated with metabolic inhibitors (2.5 μg/ml oligomycin and 1 mM 2-deoxy-D-glucose) or in the presence of metabolic inhibitors and 200 μM diazoxide (Cat#: D9035, Sigma) for 30 minutes at 37°C. The efflux solution was collected, and the cells were then lysed with 1% Triton X-100 in Ringer’s solution. Rb^+^ concentrations in both the efflux solution and cell lysate were measured using the Atomic Absorption Instrument Ion Channel Reader (ICR 8100, Aurora Biomed). The Rb^+^ efflux ratio was calculated by dividing the amount of Rb^+^ in the efflux solution by the total Rb^+^ (efflux solution plus cell lysate). The background Rb^+^ efflux ratio from untransfected cells was subtracted from other experimental readings. Data was normalized to the Rb^+^ efflux ratio of WT channels treated with metabolic inhibitors. For each experiment, technical duplicates were performed, and the average value was used as a single experimental result. Three independent transfections were performed for each condition as biological repeats.

### Inside-out Patch-clamp electrophysiology recordings

For electrophysiology, Cosm6 cells were transfected as described above, with the addition of GFP cDNA to identify the transfected cells. Recordings were performed 48– 72 hours after transfection at room temperature. The standard bath (intracellular) and pipette (extracellular) solution, referred to as K-INT/EDTA (140 mM KCl, 10 mM HEPES, 1 mM EGTA, 1mM EDTA, pH 7.2-7.4). Micro-capillary tubes (Kimble Chase Life Science) were used to pull micropipettes using a horizontal puller (Sutter Instrument Co., Novato, CA), with electrode resistance typically between 2–3 megaohms when filled with K-INT/EDTA solution. Inside-out patches were voltage-clamped using an Axopatch 1D amplifier (Axon Inc., Foster City, CA). Diazoxide, ATP (Cat#: A8937, Sigma), and ADP (Cat#: A5285, Sigma) potassium salts were used for the experiments. MgCl₂ was added to the ATP-, ADP-, or diazoxide-containing K-INT solutions to maintain a free Mg²^+^ concentration of approximately 1 mM(46). Currents were recorded at a membrane potential of -50 mV (pipette voltage set at +50 mV). Data acquisition and analysis were carried out using pCLAMP10 software (Axon Instruments)

### Statistical analysis

All data were analyzed using GraphPad Prism 8. Results are expressed as mean ± standard error of the mean. Differences were tested using one-way ANOVA, followed by a Tukey’s post hoc test for multiple comparisons. Statistical significance was set at p <0.05.

### Data availability

All data are contained within the manuscript.

## Acknowledgment

We acknowledge support from the National Institutes of Health grants R01GM145784 and R35GM161749 (to SLS), an American Heart Association Postdoctoral Scholar Award (25POST1373218; to YYK), the Cystic Fibrosis Foundation (LUECK20GO; to JDL), and the National Institutes of Health grant R01HL153988-01A1 (to JDL).

