## supplement figures for "Rescue of K_ATP_ channel nonsense mutations using anticodon-edited transfer RNAs"

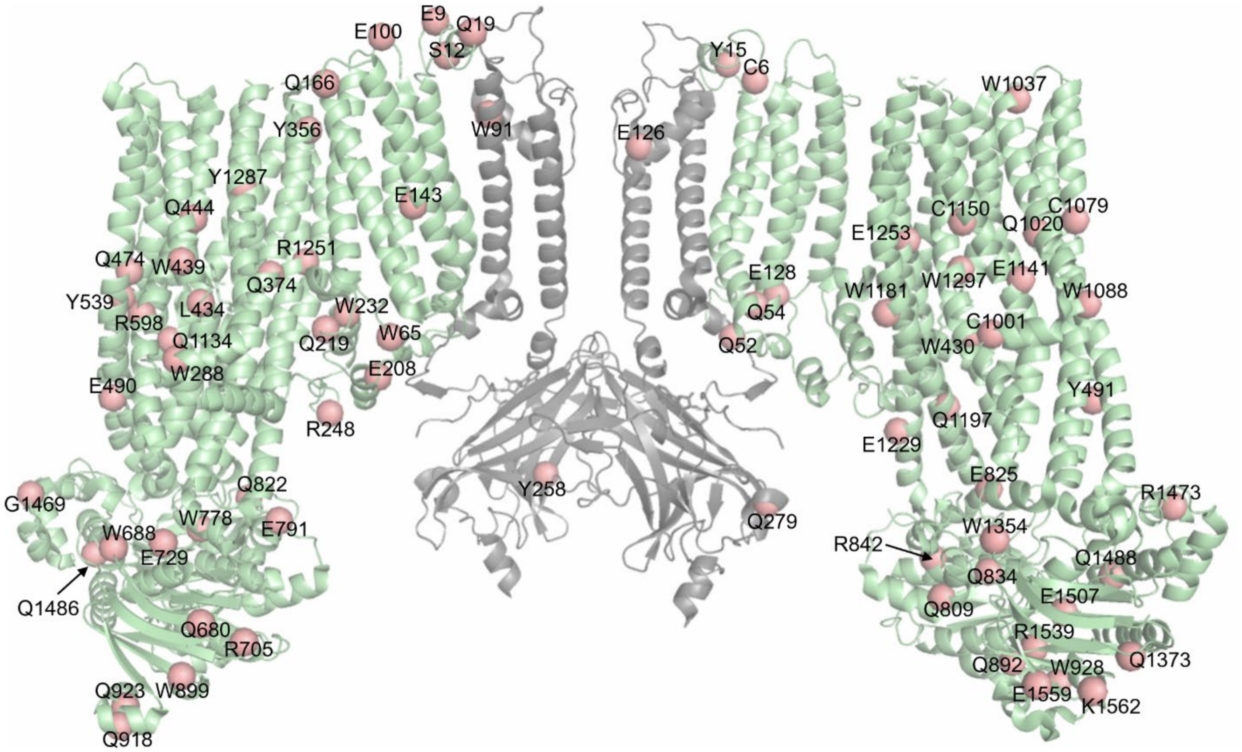


**Figure S1. Positions of $K_{ATP}$ channel PTC mutations.** PTC mutations are shown as the red spheres on the $K_{ATP}$ channel cryo-EM structure (PDB ID: 6BAA). For clarity, only two SUR1 subunits (in green) and two Kir6.2 subunits (in gray) are shown. Note, the following PTC mutations are in the unresolved regions of the channel and have not been modeled on the structure: SUR1-K329, Q335, Y344, E654, Q664, E757, R934, Q935, E946, Q954, E982, E995, W998, R999, Q1106, E1324, Y1326, E1327, W1339, and Q1342; and Kir6.2-Y12.

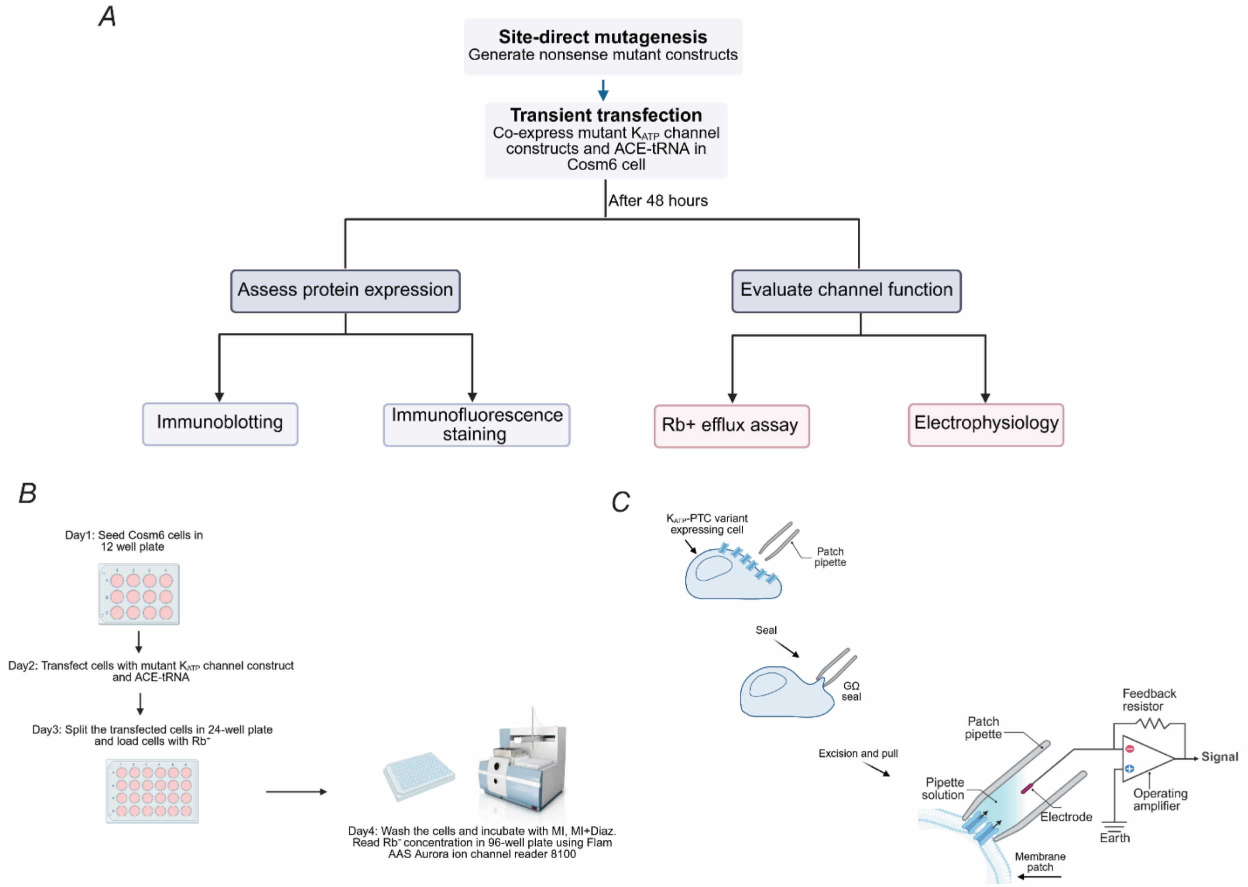


**Figure S2. Experimental schematics.** (*A*) Overview of the experimental workflow used in this study. (*B)* The experimental workflow of the $Rb^+$ efflux assay used to test the functional rescue of $K_{ATP}$-PTCs variants by ACE-tRNAs. Metabolic inhibition (MI) and metabolic inhibition+Diazoxide (MI+Diaz) were used to open $K_{ATP}$ channels, allowing $Rb^+$ efflux to be measured. (*C)* Schematic of inside-out patch-clamp electrophysiology recording. After a seal is formed between the patch pipette and the cell membrane, the membrane patch is excised, exposing the intracellular side of the membrane to the bath solution.

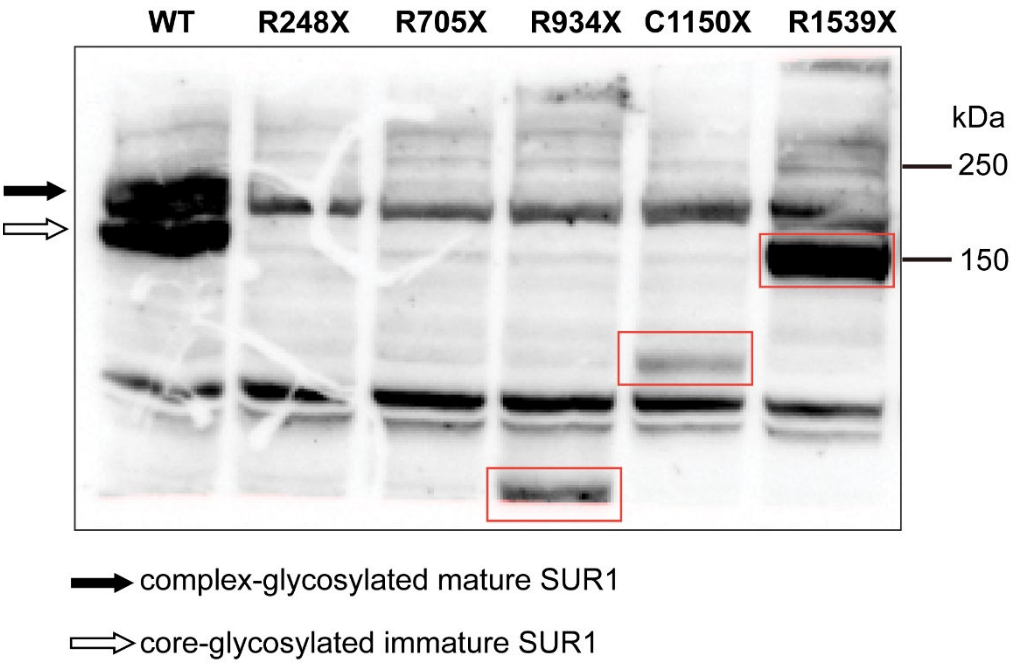


**Figure S3. Expression of truncated SUR1 proteins.** Immunoblotting with an anti-FLAG antibody shows the expression of the truncated SUR1 protein (red box).

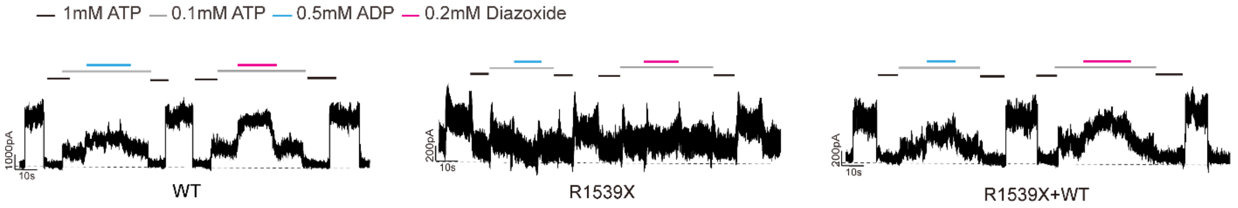


**Figure S4. Functional characterization of the heterozygous SUR1-R1539X channel.** Representative electrophysiology traces of the WT, R1539X, and R1539X+WT channels. R1539X mutant channels exhibited detectable basal currents but failed to respond to MgADP and Diazoxide. In contrast, heterozygous R1539X+WT channels showed MgADP and Diazoxide responses similar to WT channels. Holding potential: −50 mV; upward deflections indicate inward currents; black dashed lines indicate the 0 current baseline in 1 mM ATP.

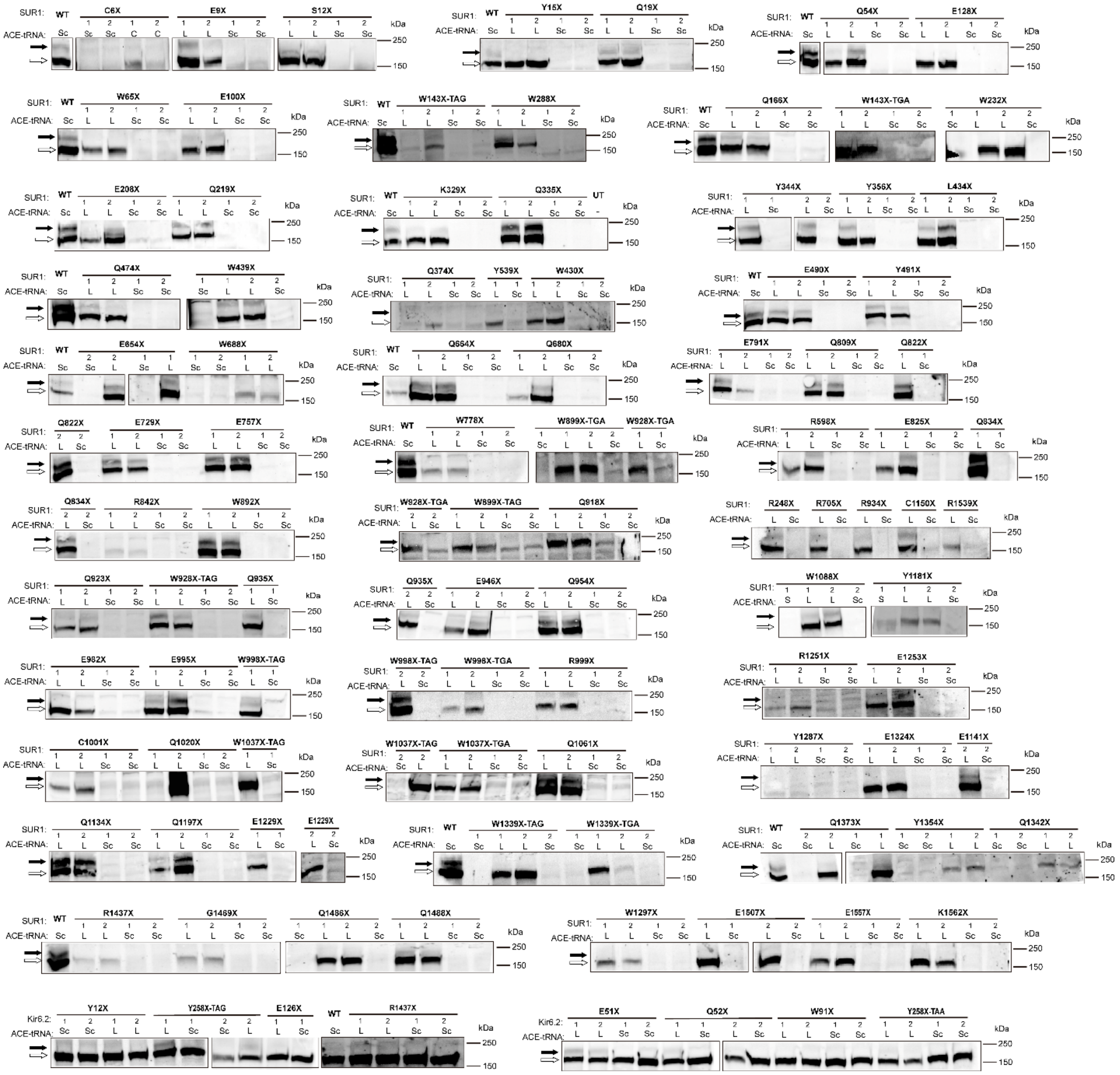

**Figure S5. Representative immunoblotting results of $K_{ATP}$ channels PTC variants screened for opt.ACE-tRNA$^{Leu}$ suppression.** Representative immunoblotting images are shown for all $K_{ATP}$ channel PTC variants tested in this study. Two independent mutant plasmid clones were generated for each variant. SUR1 expression was detected with ACE-tRNA$^{Sc}$ or opt.ACE-tRNA$^{Leu}$.

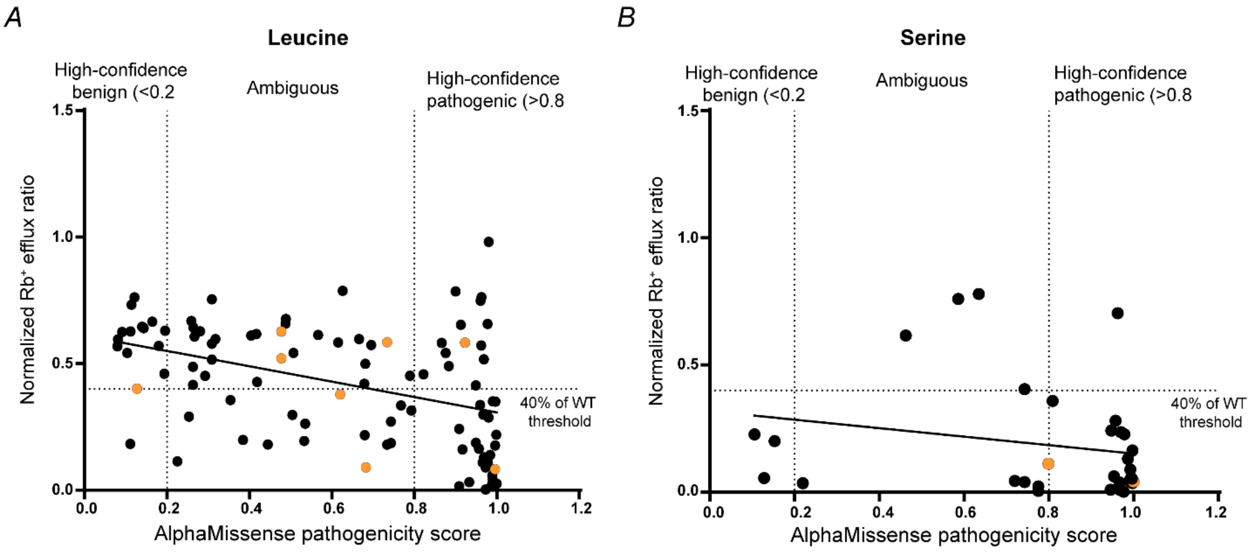


**Figure S6. Comparison of $K_{ATP}$-PTC variant functions with AlphaMissense-predicted pathogenicity scores.** *(A)* The function of the 100 $K_{ATP}$-PTC variants upon rescue by opt.ACE-tRNA$^{Leu}$ ($Rb^+$ efflux ratio under metabolic inhibition conditions) from Figure 3 is plotted against the AlphaMissense-predicted pathogenicity score for the corresponding leucine substitution. The coefficient of determination ($R^2$) for the correlation line was 0.1656. *(B)* The function of the 33 $K_{ATP}$-PTC variants ($Rb^+$ efflux ratio under metabolic inhibition condition) from Figure 4 is plotted against the pathogenicity score predicted for the serine incorporation from AlphaMissense (https://alphamissense.hegelab.org/). The coefficient of determination ($R^2$) for the correlation line was 0.03910. Black dots, SUR1-PTC variants; orange dots, Kir6.2-PTC variants.

**Table S1. PTCs identified in *ABCC8* with corresponding nucleotide changes, and the ACE-tRNA anticodon used for suppression.**

| SUR1-PTCs | | | | |
|---|---|---|---|---|
| Nucleotide | Change | Codon | amino acid change | tRNA |
| 18 | C>A | 6 | Cys-STP(TGA) | UGA |
| 25 | G>T | 9 | Glu-STP(TAG) | UAG |
| 35 | C>A | 12 | Ser-STP(TAG) | UAG |
| 45 | C>G | 15 | Tyr-STP(TAG) | UAG |
| 55 | C>T | 19 | Gln-STP(TAG) | UAG |
| 160 | C>T | 54 | Gln-STP(TAG) | UAG |
| 195 | G>A | 65 | Trp-STP(TGA) | UGA |
| 298 | G>T | 100 | Glu-STP(TAA) | UAA |
| 382 | G-T | 128 | Glu-STP(TAG) | UAG |
| 428 | G>A | 143 | Trp-STP(TAG) | UAG |
| 429 | G>A | 143 | Trp-STP(TGA) | UGA |
| 496 | C>T | 166 | Gln-STP(TAG) | UAG |
| 622 | G>T | 208 | Glu-STP(TAG) | UAG |
| 655 | C>T | 219 | Gln-STP(TAG) | UAG |
| 695 | G>A | 232 | Trp-STP(TAG) | UAG |
| 742 | C>T | 248 | Arg-STP(TGA) | UGA |
| 863 | G-A | 288 | Trp-STP(TAG) | UAG |
| 985 | A>T | 329 | Lys-STP(TAG) | UAG |
| 1003 | C>T | 335 | Gln-STP(TAG) | UAG |
| 1032 | C>G | 344 | Tyr-STP(TAG) | UAG |
| 1068 | C>G | 356 | Tyr-STP(TAG) | UAG |
| 1120 | C>T | 374 | Gln-STP(TAA) | UAA |
| 1290 | G>A | 430 | Trp-STP(TGA) | UGA |
| 1301 | T>A | 434 | Leu-STP(TAG) | UAG |
| 1317 | G>A | 439 | Trp-STP(TGA) | UGA |
| 1330 | C>T | 444 | Gln-STP(TAG) | UAG |
| 1420 | C>T | 474 | Gln-STP(TAG) | UAG |
| 1468 | G>T | 490 | Glu-STP(TAG) | UAG |
| 1473 | T>G | 491 | Tyr-STP(TAG) | UAG |
| 1617 | T>A | 539 | Tyr-STP(TAA) | UAA |
| 1792 | C>T | 598 | Arg-STP(TGA) | UGA |
| 1960 | G>T | 654 | Glu-STP(TAG) | UAG |
| 1990 | C>T | 664 | Gln-STP(TAG) | UAG |
| 2038 | C>T | 680 | Gln-STP(TAG) | UAG |
| 2064 | G>A | 688 | Trp-STP(TGA) | UGA |
| 2113 | C>T | 705 | Arg-STP(TGA) | UGA |

| Nucleotide | Change | Codon | amino acid change | tRNA |
|---|---|---|---|---|
| 2185 | G>T | 729 | Glu-STP(TAG) | UAG |
| 2269 | G>T | 757 | Glu-STP(TAG) | UAG |
| 2334 | G>A | 778 | Trp-STP(TGA) | UGA |
| 2371 | G>T | 791 | Glu-STP(TAG) | UAG |
| 2425 | C>T | 809 | Gln-STP(TAG) | UAG |
| 2464 | C>T | 822 | Gln-STP(TAG) | UAG |
| 2473 | G>T | 825 | Glu-STP(TAA) | UAA |
| 2500 | C>T | 834 | Gln-STP(TAA) | UAA |
| 2524 | C>T | 842 | Arg-STP(TGA) | UGA |
| 2674 | C>T | 892 | Gln-STP(TAG) | UAG |
| 2696 | G>A | 899 | Trp-STP(TAG) | UAG |
| 2697 | G>A | 899 | Trp-STP(TGA) | UGA |
| 2752 | C>T | 918 | Gln-STP(TAG) | UAG |
| 2767 | C>T | 923 | Gln-STP(TAG) | UAG |
| 2783 | G>A | 928 | Trp-STP(TAG) | UAG |
| 2784 | G>A | 928 | Trp-STP(TGA) | UGA |
| 2800 | C>T | 934 | Arg-STP(TGA) | UGA |
| 2803 | C>T | 935 | Gln-STP(TAG) | UAG |
| 2836 | G>T | 946 | Glu-STP(TAG) | UAG |
| 2860 | C>T | 954 | Gln-STP(TAG) | UAG |
| 2944 | G-T | 982 | Glu-STP(TAG) | UAG |
| 2983 | G-T | 995 | Glu-STP(TAG) | UAG |
| 2993 | G>A | 998 | Trp-STP(TAG) | UAG |
| 2994 | G>A | 998 | Trp-STP(TGA) | UGA |
| 2995 | C>T | 999 | Arg-STP(TGA) | UGA |
| 3003 | C>A | 1001 | Cys-STP(TGA) | UGA |
| 3058 | C-T | 1020 | Gln-STP(TAG) | UAG |
| 3110 | G>A | 1037 | Trp-STP(TAG) | UAG |
| 3111 | G>A | 1037 | Trp-STP(TGA) | UGA |
| 3181 | C>T | 1061 | Gln-STP(TAG) | UAG |
| 3237 | C>A | 1079 | Cys-STP(TGA) | UGA |
| 3263 | G>A | 1088 | Trp-STP(TAG) | UAG |
| 3400 | C>T | 1134 | Gln-STP(TAG) | UAG |
| 3421 | G>T | 1141 | Glu-STP(TAG) | UAG |
| 3450 | T>A | 1150 | Cys-STP(TGA) | UGA |
| 3543 | C>A | 1181 | Tyr-STP(TAA) | UAA |
| 3589 | C>T | 1197 | Gln-STP(TAG) | UAG |
| 3685 | G>T | 1229 | Glu-STP(TAA) | UAA |
| 3751 | C>T | 1251 | Arg-STP(TGA) | UGA |

| Nucleotide | Change | Codon | amino acid change | tRNA |
|---|---|---|---|---|
| 3757 | G>T | 1253 | Glu-STP(TAG) | UAG |
| 3861 | C>A | 1287 | Tyr-STP(TAA) | UAA |
| 3891 | G>A | 1297 | Trp-STP(TGA) | UGA |
| 3970 | G>T | 1324 | Glu-STP(TAG) | UAG |
| 4016 | G>A | 1339 | Trp-STP(TGA) | UGA |
| 4017 | G>A | 1339 | Trp-STP(TAG) | UAG |
| 4024 | C>T | 1342 | Gln-STP(TAA) | UAA |
| 4062 | c>a | 1354 | Tyr-STP(TAA) | UAA |
| 4117 | C>T | 1373 | Gln-STP(TAG) | UAG |
| 4309 | C>T | 1437 | Arg-STP(TGA) | UGA |
| 4405 | G>T | 1469 | Gly-STP(TGA) | UGA |
| 4456 | C>T | 1486 | Gln-STP(TAG) | UAG |
| 4462 | C>T | 1488 | Gln-STP(TAG) | UAG |
| 4519 | G>T | 1507 | Glu-STP(TAG) | UAG |
| 4615 | C>T | 1539 | Arg-STP(TGA) | UGA |
| 4675 | G>T | 1559 | Glu-STP(TAG) | UAG |
| 4684 | A>T | 1562 | Lys-STP(TAG) | UAG |

**Table S2. PTCs identified in *KCNJ11* with corresponding nucleotide changes, and the ACE-tRNA anticodon used for suppression.**

| Kir6.2-PTCs | | | | |
|---|---|---|---|---|
| Nucleotide | Change | Codon | amino acid change | tRNA |
| 36 | C-A | 12 | Tyr-STP(TAA) | UAA |
| 151 | G>T | 51 | Glu-STP(TAG) | UAG |
| 154 | C>T | 52 | Gln-STP(TAG) | UAG |
| 272 | G-A | 91 | Trp-STP(TAG) | UAG |
| 376 | G>T | 126 | Glu-STP(TAG) | UAG |
| 774 | C-G | 258 | Tyr-STP(TAG) | UAG |
| 774 | C-A | 258 | Tyr-STP(TAA) | UAA |
| 835 | C>T | 279 | Gln-STP(TAG) | UAG |

1 agaggcgcgg caggcgcgcg gagccagcgg agccagctga gcccgagccc agcccgcgcc
61 cgcgccgcca tgcccctggc cttctgcggc agcgagaacc actcggccgc ctaccgggtg
121 gaccaggggg tcctcaacaa cggctgcttt gtggacgcgc tcaacgtggt gccgcacgtc
181 ttcctactct tcatcacctt ccccatcctc ttcattggat ggggaagtca gagctccaag
241 gtgcacatcc accacagcac atggcttcat ttccctgggc acaacctgcg gtggatcctg
301 accttcatgc tgctcttcgt cctggtgtgt gagattgcag agggcatcct gtctgatggg
361 gtgaccgaat cccaccatct gcacctgtac atgccagccg ggatggcgtt catggctgct
421 gtcacctccg tggtctacta tcacaacatc gagacttcca acttccccaa gctgctaatt
481 gccctgctgg tgtattggac cctggccttc atcaccaaga ccatcaagtt tgtcaagttc
541 ttggaccacg ccatcggctt ctcgcagcta cgcttctgcc tcacagggct gctggtgatc
601 ctctatggga tgctgctcct cgtggaggtc aatgtcatca gggtgaggag atacatcttc
661 ttcaagacac cgagggaggt gaagcctccc gaggacctgc aagacctggg ggtacgcttc
721 ctgcagccct tcgtgaatct gctgtccaaa ggcacctact ggtggatgaa cgccttcatc
781 aagactgccc acaagaagcc catcgacttg cgagccatcg ggaagctgcc catcgccatg
841 agggccctca ccaactacca acggctctgc gaggcctttg acgcccaggt gcggaaggac
901 attcagggca ctcaaggtgc ccgggccatc tggcaggcac tcagccatgc cttcgggagg
961 cgcctggtcc tcagcagcac tttccgcatc ttggccgacc tgctgggctt cgccgggcca
1021 ctgtgcatct ttgggatcgt ggaccacctt gggaaggaga acgacgtctt ccagcccaag
1081 acacaatttc tcggggttta ctttgtctca tcccaagagt tccttgccaa tgcctacgtc
1141 ttagctgtgc ttctgttcct tgccctccta ctgcaaagga catttctgca agcatcctac
1201 tatgtggcca ttgaaactgg aattaacttg agaggagcaa tacagaccaa gatttacaat
1261 aaaattatgc acctgtccac ctccaacctg tccatgggag aaatgactgc tggacagatc
1321 tgtaatctgg ttgccatcga caccaatcag ctcatgtggt ttttcttctt gtgcccaaac
1381 ctctgggcta tgccagtaca gatcattgtg ggtgtgattc tcctctacta catactcgga
1441 gtcagtgcct taattggagc agctgtcatc attctactgg ctcctgtcca gtacttcgtg
1501 gccaccaagc tgtctcaggc ccagcggagc acactggagt attccaatga gcggctgaag
1561 cagaccaacg agatgctccg cggcatcaag ctgctgaagc tgtacgcctg ggagaacatc
1621 ttccgcacgc gggtggagac gacccgcagg aaggagatga ccagcctcag ggcctttgcc
1681 atctatacct ccatctccat tttcatgaac acggccatcc ccattgcagc tgtcctcata
1741 actttcgtgg gccacgtcag cttcttcaaa gaggccgact tctcgccctc cgtggccttt
1801 gcctccctct ccctcttcca tatcttggtc acaccgctgt tcctgctgtc cagtgtggtc
1861 cgatctaccg tcaaagctct agtgagcgtg caaaagctaa gcgagttcct gtccagtgca
1921 gagatccgtg aggagcagtg tgccccccat gagcccacac ctcagggccc agccagcaag
1981 taccaggcgg tgcccctcag ggttgtgaac cgcaagcgtc cagcccggga ggattgtcgg
2041 ggcctcaccg gcccactgca gagcctggtc cccagtgcag atggcgatgc tgacaactgc
2101 tgtgtccaga tcatgggagg ctacttcacg tggaccccag atggaatccc cacactgtcc
2161 aacatcacca ttcgtatccc ccgaggccag ctgactatga tcgtggggca ggtgggctgc
2221 ggcaagtcct cgctccttct agccgcactg gggggagatgc agaaggtctc aggggctgtc
2281 ttctggagca gcagccttcc tgacagcgag ataggagagg accccagccc agagcgggag
2341 acagcgaccg acttggatat caggaagaga ggccccgtgg cctatgcttc gcagaaacca
2401 tggctgctaa atgccactgt ggaggagaac atcatctttg agagtccctt caacaaacaa
2461 cggtacaaga tggtcattga agcctgctct ctgcagccag acatcgacat cctgccccat
2521 ggagaccaga cccagattgg ggaacggggc atcaacctgt ctggtggtca acgccagcga
2581 atcagtgtgg cccgagccct ctaccagcac gccaacgttg tcttcttgga tgaccccttc
2641 tcagctctgg atatccatct gagtgaccac ttaatgcagg ccggcatcct tgagctgctc
2701 cgggacgaca agaggacagt ggtcttagtg acccacaagc tacagtacct gccccatgca
2761 gactggatca ttgccatgaa ggatggcacc atccagaggg agggtaccct caaggacttc
2821 cagaggtctg aatgccagct ctttgagcac tggaagaccc tcatgaaccg acaggaccaa
2881 gagctggaga aggagactgt cacagagaga aaagccacag agccacccca gggcctatct
2941 cgtgccatgt cctcgaggga tggccttctg caggatgagg aagaggagga agaggaggca
3001 gctgagagcg aggaggatga caacctgtcg tccatgctgc accagcgtgc tgagatccca
3061 tggcgagcct gcgccaagta cctgtcctcc gccggcatcc tgctcctgtc gttgctggtc
3121 ttctcacagc tgctcaagca catggtcctg gtggccatcg actactggct ggccaagtgg
3181 accgacagcg ccctgaccct gacccctgca gccaggaact gctccctcag ccaggagtgc
3241 accctcgacc agactgtcta tgccatggtg ttcacggtgc tctgcagcct gggcattgtg
3301 ctgtgcctcg tcacgtctgt cactgtggag tggacagggc tgaaggtggc caagagactg
3361 caccgcagcc tgctaaaccg gatcatccta gcccccatga ggtttttttga gaccacgccc

```
3421 cttgggagca tcctgaacag attttcatct gactgtaaca ccatcgacca gcacatccca
3481 tccacgctgg agtgcctgag ccgctccacc ctgctctgtg tctcagccct ggccgtcatc
3541 tcctatgtca cacctgtgtt cctcgtggcc ctcttgcccc tggccatcgt gtgctacttc
3601 atccagaagt acttccgggt ggcgtccagg gacctgcagc agctggatga caccacccag
3661 cttccacttc tctcacactt tgccgaaacc gtagaaggac tcaccaccat ccgggccttc
3721 aggtatgagg cccggttcca gcagaagctt ctcgaataca cagactccaa caacattgct
3781 tccctcttcc tcacagctgc caacagatgg ctggaagtcc gaatggagta catcggtgca
3841 tgtgtggtgc tcatcgcagc ggtgacctcc atctccaact ccctgcacag ggagctctct
3901 gctggcctgg tgggcctggg ccttacctac gccctaatgg tctccaacta cctcaactgg
3961 atggtgagga acctggcaga catggagctc cagctggggg ctgtgaagcg catccatggg
4021 ctcctgaaaa ccgaggcaga gagctacgag ggctcctgg caccatcgct gatcccaaag
4081 aactggccag accaagggaa gatccagatc cagaacctga gcgtgcgcta cgacagctcc
4141 ctgaagccgg tgctgaagca cgtcaatgcc ctcatcgccc ctggacagaa gatcgggatc
4201 tgcggccgca ccggcagtgg gaagtcctcc ttctctcttg ccttcttccg catggtggac
4261 acgttcgaag ggcacatcat cattgatggc attgacatcg ccaaactgcc gctgcacacc
4321 ctgcgctcac gcctctccat catcctgcag gaccccgtcc tcttcagcgg caccatccga
4381 tttaacctgg accctgagag gaagtgctca gatagcacac tgtgggaggc cctggaaatc
4441 gcccagctga agctggtggt gaaggcactg ccaggaggcc tcgatgccat catcacagaa
4501 ggcggggaga atttcagcca gggacagagg cagctgttct gcctggcccg ggccttcgtg
4561 aggaagacca gcatcttcat catggacgag gccacggctt ccattgacat ggccacggaa
4621 aacatcctcc aaaaggtggt gatgacagcc ttcgcagacc gcactgtggt caccatcgcg
4681 catcgagtgc acaccatcct gagtgcagac ctggtgatcg tcctgaagcg ggtgccatc
4741 cttgagttcg ataagccaga gaagctgctc agccggaagg acagcgtctt cgcctccttc
4801 gtccgtgcag acaagtgacc tgccagagcc caagtgccat cccacattcg gaccctgccc
4861 ataccccctgc ctgggttttc taactgtaaa tcacttgtaa ataaatagat ttgattattt
4921 ccta
```

File S1. The complete DNA sequence of *ABCC8* used in this study. Ref:[NM_001287174.3]

```
   1 ggcagggttg ggggctccct aggcgccagg caggtgggct caagggtgag gctgtttttt
  61 ttgttttgtt ttttgttttt gagacggagt ctcgctctgt cgcccaggct ggagtgcagt
 121 ggcgtgatct tggctcactg caacctccgc ctctcgggtt caagcgattc tcctgcctca
 181 gcttcctgag tagctgggat tacaggtgcg caccaccatg cccggctaac ttttgtattt
 241 ttagtagaga tggggtttca ccatgttggt caggctggtc tcgaactcct gacctagtga
 301 tctgccctcc tcagcctccc aacgtactgg gattacaggc gtgagccacc gcgcccggcc
 361 tgaggctggt attaagaagt gaagtgggac ccaggtggag gtaaggaaga gtctggtggg
 421 gagttatctc agaagtgagg ccagcacagg ctgagtgcag ccccagggtg agaaggtgcc
 481 caccgagagg actctgcagt gaggccctag gccacgtccg aggggtgcct ccgatggggg
 541 aagcccctcc ctgggggtca ccggagccat gctgtcccgc aagggcatca tccccgagga
 601 atacgtgctg acacgcctgg cagaggaccc tgccaagccc aggtaccgtg cccgccagcg
 661 gagggcccgc tttgtgtcca agaaaggcaa ctgcaacgtg gcccacaaga acatccggga
 721 gcagggccgc ttcctgcagg acgtgttcac cacgctggtg gacctcaagt ggccacacac
 781 attgctcatc ttcaccatgt ccttcctgtg cagctggctg ctcttcgcca tggcctggtg
 841 gctcatcgcc ttcgcccacg gtgacctggc cccagcgag ggcactgctg agccctgtgt
 901 caccagcatc cactccttct cgtctgcctt ccttttctcc attgaggtcc aagtgactat
 961 tggctttggg gggcgcatgg tgactgagga gtgcccactg gccatcctga tcctcatcgt
1021 gcagaacatc gtggggctca tgatcaacgc catcatgctt ggctgcatct tcatgaagac
1081 tgcccaagcc caccgcaggg ctgagaccct catcttcagc aagcatgcgg tgatcgccct
1141 gcgccacggc cgcctctgct tcatgctacg tgtgggtgac ctccgcaaga gcatgatcat
1201 cagcgccacc atccacatgc aggtggtacg caagaccacc agccccgagg gcgaggtggt
1261 gcccctccac caggtggaca tccccatgga gaacggcgtg ggtggcaaca gcatcttcct
1321 ggtggccccg ctgatcatct accatgtcat tgatgccaac agcccactct acgacctggc
1381 acccagcgac ctgcaccacc accaggacct cgagatcatc gtcatcctgg aaggcgtggt
1441 ggaaaccacg ggcatcacca cccaggcccg cacctcctac ctggccgatg agatcctgtg
1501 gggccagcgc tttgtgccca ttgtagctga ggaggacgga cgttactctg tggactactc
1561 caagtttggc aacaccgtca aagtgcccac accactctgc acggcccgcc agcttgatga
1621 ggaccacagc ctactggaag ctctgaccct cgcctcagcc cgcgggcccc tgcgcaagcg
1681 cagcgtgccc atggccaagg ccaagcccaa gttcagcatc tctccagatt ccctgtcctg
1741 agccatggtc tctcgggccc cccacacgcg tgtgtacaca cggaccatgt ggtatgtagc
1801 ccggccaggg cctggtgtga ggctgggcca gcctcagctc agcctccccc tgctgctcat
1861 ccagggtgtt acaaggcact tgtcactatg ctatttctgg cctcagcagg aacctgtact
1921 gggttatttt tgtccctgct cctcccaacc caattcagga ctggctcacc cctctccccc
1981 gcccaaggct gcagaggctg tgggaggtac tgggccctag agctgtgcgt ccagccagtc
2041 ctgggtcccc acgattgacc agccacactc tgggccggtg gctggggaag aacaatcccc
2101 gagggctgct gctttgcgtc tgtggctcca agaagtgcct gtggtcaggc cccagctcta
2161 cttggtccct gaaaaagcac ctggctaagg gctgggcctg gccagcaggg agggcagttg
2221 atgagagagg gtgttcccgc tggagggttg gtgctgtgga gcctacaccg gcagggacag
2281 cctggggctg acagggctcc cctccgaggg ccagtttcag gtctggaagg ggaggaagca
2341 ggggaaggtg acctgaggag gctcggcttt gtagagcccc gctcaggcac agggaggagg
2401 agatgccagg gctcctgcct tttgccacat cggcctcgtg cagtgagggc tctgtgggct
2461 ggggctgctg cccctgccta cctcctgcct gtccccagag gctgaggaga gggggtactg
2521 tgcccaccac acatgattag gcctcagacc caactctggt cctggctcca caacagtggc
2581 tgccactcac tttgtccaga aggtggcttg gggtggata tctttgggtt gctggaaaag
2641 gtgtgggaag gttcaggatg gtgggaggga ctgaggtccc tgaggtgaag aggcccttgg
2701 tcctgacggg tttgacccgt gcctggaccc ttggagcagt gttgtgtgaa cttgcctaga
2761 actctgcctt ctccgttgtc aataaagcct ccccctcatg acctaaactc tgggcttttc
2821 ttgctgggga ggcagcaagc atgctggtgg gaagggaggc agggactggc agctgccacc
2881 cccttcaaga ggcgccatag accctagcgg ggagggcagg ggagggacgg aaggctggca
2941 cctcttccac cagttcaggg ggactttccc ctctcctgtc tcaggtggcc cagccctgtc
3001 agcctgtctg gccaactcag cctttgggca ctcaccaggc tttgcagccc tgggctctgt
3061 ctctactccc agggacctgc tggaaggctg gagtgcccag ggagaggtat agaggtgtca
3121 taggcattag tgtagtaatt ggagcactaa ctctcgagcc aactgcctgg gttcgaatcc
3181 tggctctagc tgtatgactt ttgtcaagta acttagcctc tctgtgtctc agttgcctct
3241 tctataacat ggatgctaat agtacctacc tcatagaatt gttttggaag taaatgaaaa
3301 atatgtaaaa tgctgaagtg cctggtctac agtaagtgct caataaatgt taactattgt
3361 gattgctgct gaatcagcta catgctgagg aaacggccaa acaagtgtta aaaaaaaa
```

File S2. The complete DNA sequence of *KCNJ11* used in this study. Ref:[NM_000525.3]
